# Demonstrating and engineering the *de novo* synthesis of two aromatic chemicals estragole and anethole in *Escherichia coli*

**DOI:** 10.64898/2025.12.22.695922

**Authors:** Yongkun Lv, Shuai Ding, Jinmian Chang, Ying Xiao, Shiming Gu, Anqi Zhao, Shilei Wang, Weigao Wang, Yameng Xu, Jingliang Xu, Peng Xu

**Author notes:** Corresponding Authors: Peng Xu, Jingliang Xu, and Jinmian Chang.

## Abstract

Estragole and anethole are the major flavoring and aroma components of the essential oils extracted from basil (*Ocimum basilicum* L.), anise (*Pimpinella anisum*), star anise (*Illicium verum*), and fennel (*Foeniculum vulgare*). Besides, they also exhibit bioactivities, such as antioxidation, antimicrobial, immunomodulation, inti-inflammation, and anti-diabetes. These properties make them being faced with substantial applications in the industries of food, cosmetics, pharmaceutics, and agriculture. Commercial estragole and anethole are currently obtained from extraction from their original plants, which suffers from unstable quality and resources. Microbial synthesis is a promising alternative approach to sustainably obtain estragole and anethole, but has not been achieved. In this study, we modularly redesigned the synthetic pathways based on the *Escherichia coli* chassis. Firstly, a *p*-coumaryl alcohol over-accumulator (436.28 mg/L) was engineered to facilitate the validation of the undemonstrated downstream module. *De novo* biosynthesis of estragole and anethole was achieved by stepwise analysis and BioBricks assembly. The intracellular availability of *S*-adenosyl-L-methionine (SAM) was revealed to be the key limiting factor, and was enhanced by deregulating the synthesis. Finally, the productions of estragole and anethole were improved by 246.88% and 202.44% compared with the initial strains. This study represents the first synthesis of estragole and anethole in microbial chassis, and the demonstration of microbial synthesis of propenyl phenols. Besides, it provided an alternative sustainable approach to obtain flavoring and aroma chemicals.

## 1. Introduction

The aromatic and medical plants basil (*Ocimum basilicum* L.), anise (*Pimpinella anisum*), star anise (*Illicium verum*), and fennel (*Foeniculum vulgare*) are among the most widely used culinary herbs. Besides, they have long been used as the traditional medical herbs in China, India, and part of Europe ^1^. Essential oils extracted from these herbs are famous for their flavoring and aroma properties, and have been widely used in the fields of foods, cosmetics, and pharmaceuticals ^2^. The price is predominantly determined by the chemical compositions of the essential oil, which however is largely affected by the varieties of species, culture conditions, and geographic distributions ^3^. Estragole (1-allyl-4-methoxybenzene) and anethole (1-methoxy-4-[(1*E*)-prop-1-en-1-yl]benzene) are among the major components and the primary bioactive chemicals in these essential oils ^4^. As the predominant contributors to the flavoring and aroma properties and biological activities of the essential oils, estragole and anethole are faced with potential huge market demand ^5, 6^. So far, the commercial estragole and anethole are predominantly extracted from their original plants. However, the extraction is susceptible to multiple factors as aforementioned ^3^.

Engineering microbial cell factory (MCF) is a promising alternative approach to stably obtain the plant natural products ^7^. However, the biosynthesis of estragole and anethole has not been demonstrated in microbial chassis, which hinders the development of MCF. Moreover, the reactions catalyzed by coniferyl alcohol acyltransferase (CFAT), eugenol/chavicol synthase (EGS), isoeugenol/isochavicol synthase (IGS), and allyl/propenyl phenol synthase (APS/PPS), which are involved in the proposed synthetic pathways (**Figure 1**), are scarcely studied in the microbial chassis. Commercial *p*-coumaryl acetate and isochavicol are unavailable at present. On the other hand, *O*-methylation, which is a *S*-adenosyl-L-methionine (SAM) dependent reaction, is involved in the proposed biosynthetic pathway (**Figure 1**). As the primary methyl donor, the homeostasis of SAM suffers from strict multilayer regulations ^8, 9^. The limited intracellular availability makes SAM a key limiting factor for the over-production of SAM-dependent products in the engineered MCF ^10^.

**Figure 1.**
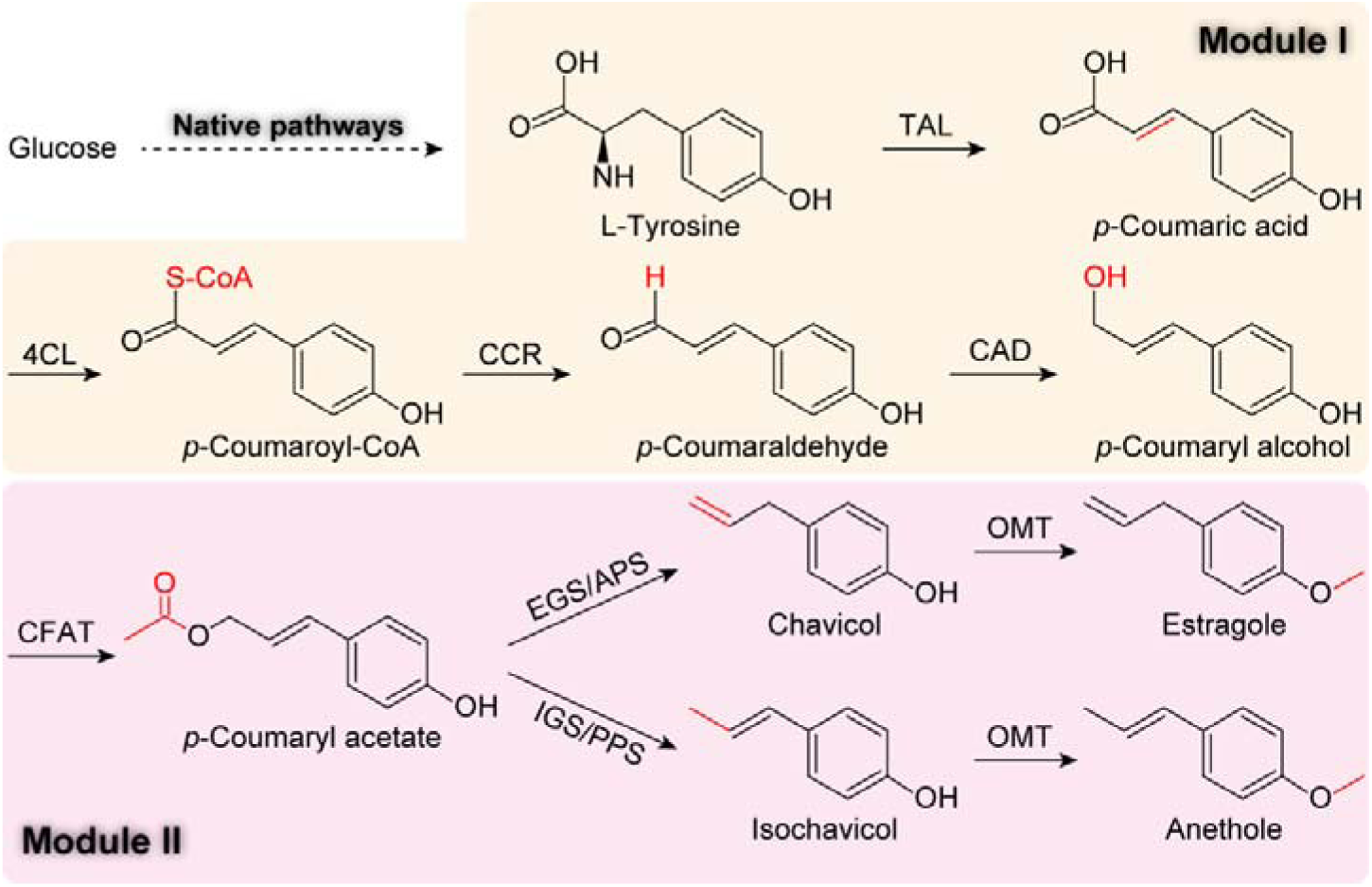
The modular design of synthetic pathways of estragole and anethole. The synthetic pathways start from L-tyrosine, which is synthesized in *E. coli* through the native pathways. The synthetic pathways of estragole and anethole were designed into two modules. Module I is composed of enzymatic reactions that have been demonstrated in the microbial chassis, while module II is composed of those have not yet been demonstrated in microbial chassis. Red indicates the changed group. TAL, tyrosine ammonia-lyase; 4CL, *p*-coumarate-CoA ligase; CCR, cinnamoyl-CoA reductase; CAD, cinnamyl-alcohol dehydrogenase; CFAT, coniferyl alcohol acyltransferase; EGS, eugenol/chavicol synthase; IGS, isoeugenol/isochavicol synthase; APS/PPS, allyl/propenyl phenol synthase; OMT, *O*-methyltransferase.

In this study, we will modularly redesign the potential synthetic pathways of estragole and anethole in the *Escherichia coli* chassis (**Figure 1**). To facilitate the validation of the unexplored module II, we will firstly engineer an over-accumulator of *p*-coumaryl alcohol, the precursor of module II. Stepwise validation will be carried out to demonstrate the biosynthesis in *E. coli* chassis. The intracellular SAM availability will also be engineered to improve the production. We will try to develop a stable and sustainable alternative approach to obtain estragole and anethole, two flavoring and aroma chemicals derived from the widely used essential oils.

## 2. Results and discussion

### 2.1 Modular design of the synthesis pathway

In *Escherichia coli* chassis, the heterologous synthetic pathways of estragole and anethole are from L-tyrosine, which can be synthesized through the native pathways (**Figure 1**). L-Tyrosine is sequentially converted into *p*-coumaric acid, *p*-coumaroyl-CoA, *p*-coumaraldehyde, *p*-coumaryl alcohol, and *p*-coumaryl acetate by the heterologous enzymes tyrosine ammonia-lyase (TAL), *p*-coumarate-CoA ligase (4CL), cinnamoyl-CoA reductase (CCR), cinnamyl-alcohol dehydrogenase (CAD) and coniferyl alcohol acyltransferase (CFAT). Thereafter, estragole and anethole are synthesized through different pathways. Eugenol/chavicol synthase (EGS) and isoeugenol/isochavicol synthase (IGS) respectively convert *p*-coumaryl acetate into chavicol and isochavicol, whose para hydroxy groups are subsequently methylated by the *O*-methyltransferase (OMT), resulting in estragole and anethole.

To facilitate the pathway building, analysis, and validation, we divided the heterologous pathways into 2 modules (**Figure 1**). Module I is from L-tyrosine to *p*-coumaryl alcohol, and this route has been validated in *E. coli* chassis ^11, 12^. Module II is from *p*-coumaryl alcohol to estragole and anethole. These routes have not yet been demonstrated in any microbial chassis. Stepwise building and validation are required on the basis of *p*-coumaryl alcohol over-accumulation.

### 2.2 Building a *p*-coumaric alcohol producer

According to the modular design, we should firstly rebuild module I and engineer a *p*-coumaryl alcohol producer. As *RgTAL* and *Pc4CL* have been analyzed and validated in our previous studies ^13^, these two genes were used directly to build the synthetic pathway herein. On this basis, we obtained 3 *p*-coumaraldehyde producing strains (CAD-01 to 03, **Table S4**) by introducing the cinnamoyl-CoA reductase (CCR) encoding genes (**Table S1 and Figure S1A**). The *p*-coumaraldehyde production was confirmed by HPLC and LC/MS analysis (**Figure 2A and B**). Among the strains, CAD-01 expressing *AtCCR* from *Arabidopsis thaliana* accumulated highest *p*-coumaraldehyde titer (9.00 mg/L) (**Figure 2C**). On the basis of *p*-coumaraldehyde production, we tried to produce *p*-coumaryl alcohol by introducing the alcohol dehydrogenase encoding genes. All the genes that are annotated as alcohol dehydrogenase or aldehyde reductase from *E. coli* and *S. cerevisiae* were tested (**Table S1**). Besides, the potential alcohol dehydrogenase encoding genes from *Yarrowia lipolytica* were also dug by using local TBLASTN (**Table S1**). Alcohol dehydrogenases from *S. cerevisiae* were used as the query ^14, 15^. The expression of these genes was confirmed by SDS-PAGE (**Figures S1B** and **C**). Introducing these genes carried by pACM4 (**Table S3**) into strain CAD-01 resulted in the *p*-coumaryl alcohol producing strains (CAL-01 to 20, **Table S4**). HPLC analysis showed that all these recombinant strains produced *p*-coumaryl alcohol, and CAL-06 expressing *ScADH6* from *S. cerevisiae* produced the highest titer (110.37 mg/L, **Figures 2D and E** and **Figures S1D-G**). However, it should be noted that the control strain CAD-01/pACM, which contained empty plasmid pACM4 without any heterologous alcohol dehydrogenase gene, also produced considerable *p*-coumaryl alcohol (83.07 mg/L, **Figures S1E and G**). We then assembled *ScADH6* with other pathway genes, resulting in the plasmid pET28a(PB)N-RgTAL-Pc4CL-AtCCR-ScADH6 (**Table S3**) and strain CAL-21 (**Table S4**). Intriguingly, the resulting strain CAL-21 produced reduced *p*-coumaryl alcohol titer compared with that of CAL-06 (**Figures 2F** and **S1F**). We speculate that moving *ScADH6* from the low copy number (10-12) pACM4 to the higher copy number (∼40) pET28a(PB)N resulted in excess alcohol dehydrogenase activity, which severely interfered with the chassis basic metabolism ^12, 16, 17^. On the contrary, removing the empty plasmid pACM4 from the control strain CAD-01/pACM4 improved *p*-coumaryl alcohol titer to 140.01 mg/L (**Figures 2F** and **S1G**). These results indicated that the activity of *E. coli* native alcohol dehydrogenases (**Table S1**) is high enough for *p*-coumaryl alcohol production, and excess overexpression could result in metabolic interference and titer reduction.

**Figure 2.**
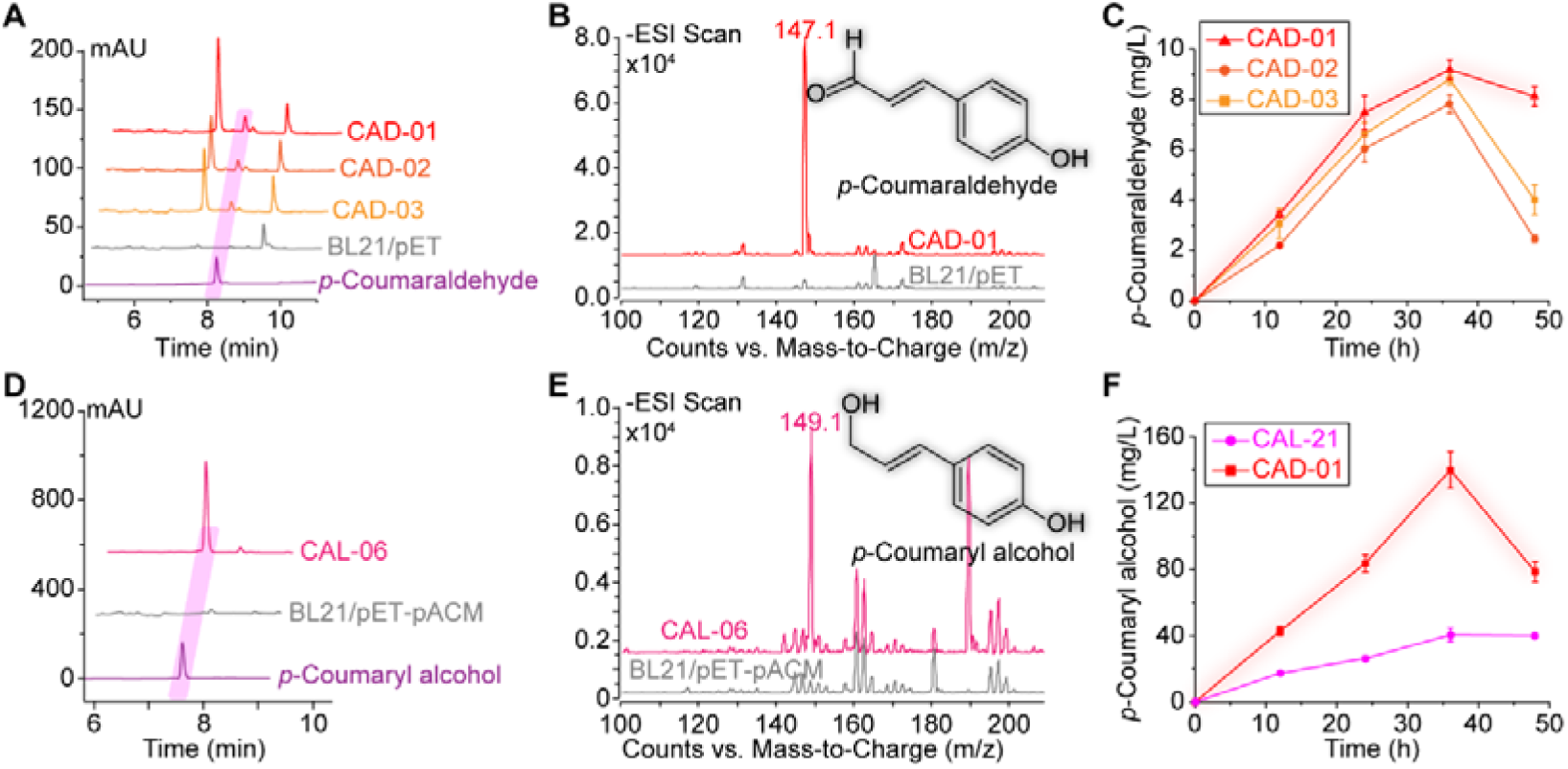
Build the *p*-coumaryl alcohol synthesis pathway in *E. coli*. **(A)** HPLC analysis of the *p*-coumaraldehyde production. **(B)** LC/MS analysis of the *p*-coumaraldehyde production. The molecular weight of *p*-coumaraldehyde is 148.16 g/mol. The mass-to-charge 147.1 m/z in negative mode indicates the production of *p*-coumaraldehyde. **(C)** *p*-Coumaraldehyde titers produced by strains overexpressing CCR encoding genes from different organisms. **(D)** HPLC analysis of the *p*-coumaryl alcohol production. **(E)** LC/MS analysis of the *p*-coumaryl alcohol production. The molecular weight of *p*-coumaryl alcohol is 150.17 g/mol. The mass-to-charge 149.1 m/z in negative mode indicates the production of *p*-coumaryl alcohol. **(F)** The *p*-coumaryl alcohol titers produced by the strains that overexpress ScADH6 (CAL-06) and that only contains native alcohol dehydrogenases (CAD-01). BL21/pET and BL21/pET-pACM containing blank plasmid(s) were used as the control sets.

### 2.3 Engineering a *p*-coumaric alcohol over-accumulating chassis

Despite the achieving of *p*-coumaryl alcohol production, we tried to further improve the accumulation of *p*-coumaryl alcohol. As module II has not yet been validated in microbial chassis, the over-accumulation of *p*-coumaryl alcohol, which serves as the substrate of module II, will facilitate the stepwise demonstration.

The biosynthesis of aromatic amino acids is tightly regulated on both the transcriptional and post-transcriptional levels, thus hinders the over-accumulation of aromatic amino acids and their derivative (**Figure 3A**).

**Figure 3.**
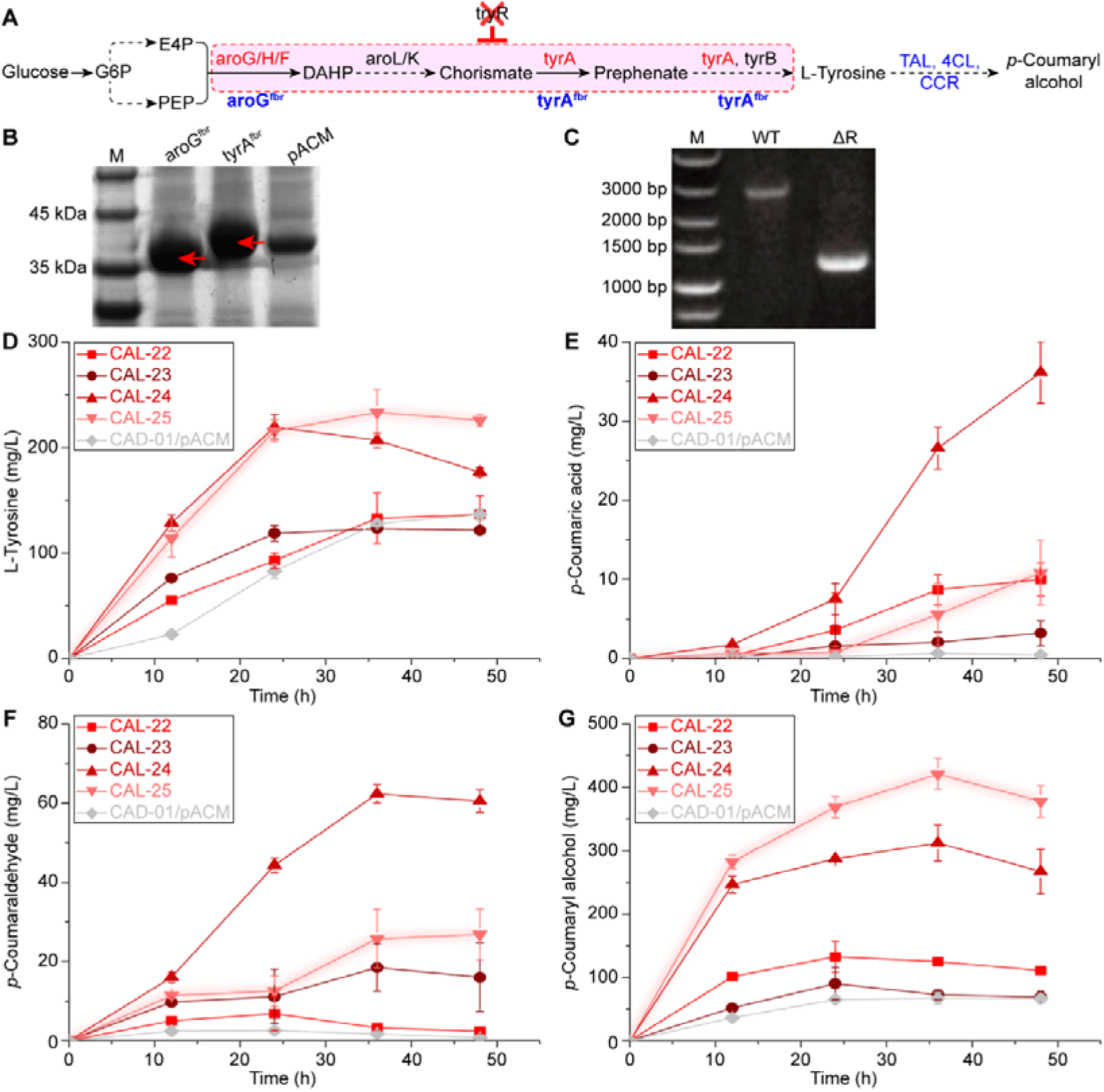
Engineering *p*-coumaryl alcohol over-accumulation by removing feedback regulation. **(A)** The synthesis pathway and regulation of *p*-coumaryl alcohol. Solid lines indicate single-step reactions. Dash lines indicate multiple-step reactions. Red color indicates that the enzyme is feedback inhibited. The genes in the dash red box are feedback repressed through tyrR. Red cross indicates gene deletion, while blue color indicates gene over-expression. **(B)** SDS-PAGE analysis of the over-expression of aroG^fbr^ and tyrA^fbr^. Molecular weights of recombinant proteins were as follows: aroG^fbr^ 41.42 kDa, tyrA^fbr^ 45.47 kDa. Red arrows indicate the specific bands. pACM, referring to *E. coli* BL21(DE3) containing empty plasmid pACM4, was used as blank control. Other lanes were *E. coli* BL21(DE3) overexpressing corresponding genes with pACM4. **(C)** Diagnostic PCR analysis of *tyrR* gene knockout. The specific bands of wild type and mutant were 2714-bp and 1172-bp. WT refers to wild type *E. coli* BL21(DE3). **(D-G)** The titers of L-tyrosine, *p*-coumaric acid, *p*-coumaraldehyde, and *p*-coumaryl alcohol accumulated by the engineered strains.

3-Deoxy-D-arabino-heptulosonate-7-phosphate (DAHP) synthase catalyzes the first committed step of the aromatic amino acid biosynthesis. *E. coli* contains three DAHP synthase isomers (aroG, aroH, and aroF), each of which suffers from feedback inhibition by L-phenylalanine, L-tryptophan, and L-tyrosine. The D146N mutant of aroG has been reported to exhibit the feedback inhibition resistant (fbr) property ^18, 19^. The chorismate mutase/prephenate dehydrogenase (tyrA), which catalyzes two sequential steps from chorismate to *p*-hydroxyphenylpyruvate, is feedback inhibited by L-tyrosine. Combinatorial mutations of M53I and A354V also released tyrR from feedback inhibition ^19-21^. We tried to achieve the over-accumulation of *p*-coumaryl alcohol by over-expressing these feedback-inhibition resistant mutants (aroG^fbr^ and tyrA^fbr^). When aroG^fbr^ and tyrA^fbr^ were overexpressed individually (CAL-22 and CAL-23, **Table S4**), only moderate improvements were observed on the titers of *p*-coumaric acid, *p*-coumaraldehyde, and *p*-coumaryl alcohol (**Figures 3B** and **3D-G**). However, when aroG^fbr^ and tyrA^fbr^ were over-expressed combinatorially (CAL-24, **Table S4**), dramatic improvements were observed on the titers of L-tyrosine, *p*-coumaric acid, *p*-coumaraldehyde, and *p*-coumaryl alcohol. The *p*-coumaryl alcohol titer reached 311.98 mg/L, representing a 362.95% improvement compared with the control strain CAD-01/pACM (**Figures 3B** and **3D-G**).

Besides the post-transcriptional feedback inhibition on enzymes, aromatic amino acid biosynthesis also suffers from transcription regulations. In *E. coli*, the biosynthesis of L-tyrosine is primarily regulated by the transcription regulator tyrR. In the presence of aromatic amino acid, the transcription of multiple genes of the synthetic pathway (such as *aroG/H/F*, *aroL*, *tyrA*, and *tyrB*) are repressed through tyrR (**Figure 3A**) ^22^. This feedback repression could be eliminated by deleting tyrR ^21, 23^. In this study, knocking out *tyrR* gene (**Figure 3C**) further improved the *p*-coumaryl alcohol titer to 421.17 mg/L, which represents a further improvement of 35.00% compared with CAL-24 (**Figure 3G**). Interestingly, the improvement of *p*-coumaryl alcohol titer was accompanied with the decrease of *p*-coumaric acid and *p*-coumaraldehyde accumulations (**Figures 3E-G**). This indicates that deleting tyrR promoted the conversion from *p*-coumaric acid and *p*-coumaraldehyde into *p*-coumaryl alcohol. As described above, the final step of *p*-coumaryl alcohol synthesis was catalyzed by the *E. coli* native alcohol dehydrogenase(s) in the engineered strains. It can be deduced that these genes might also suffer from the transcription regulation by transcription factor tyrR, which remains uncovered mechanism ^22, 24^.

L-Phenylalanine and L-tryptophan compete precursors with *p*-coumaryl alcohol. We also tried to block these competing pathways by removing chorismate mutase/prephenate dehydratase (encoded by *pheA*) and anthranilate synthase component I (encoded by *trpE*), which catalyze the first steps of the branched pathways leading to the synthesis of L-phenylalanine and L-tryptophan (**Figure 4A**). The gene knockout was confirmed using diagnostic PCR analysis (**Figure S2**).

**Figure 4.**
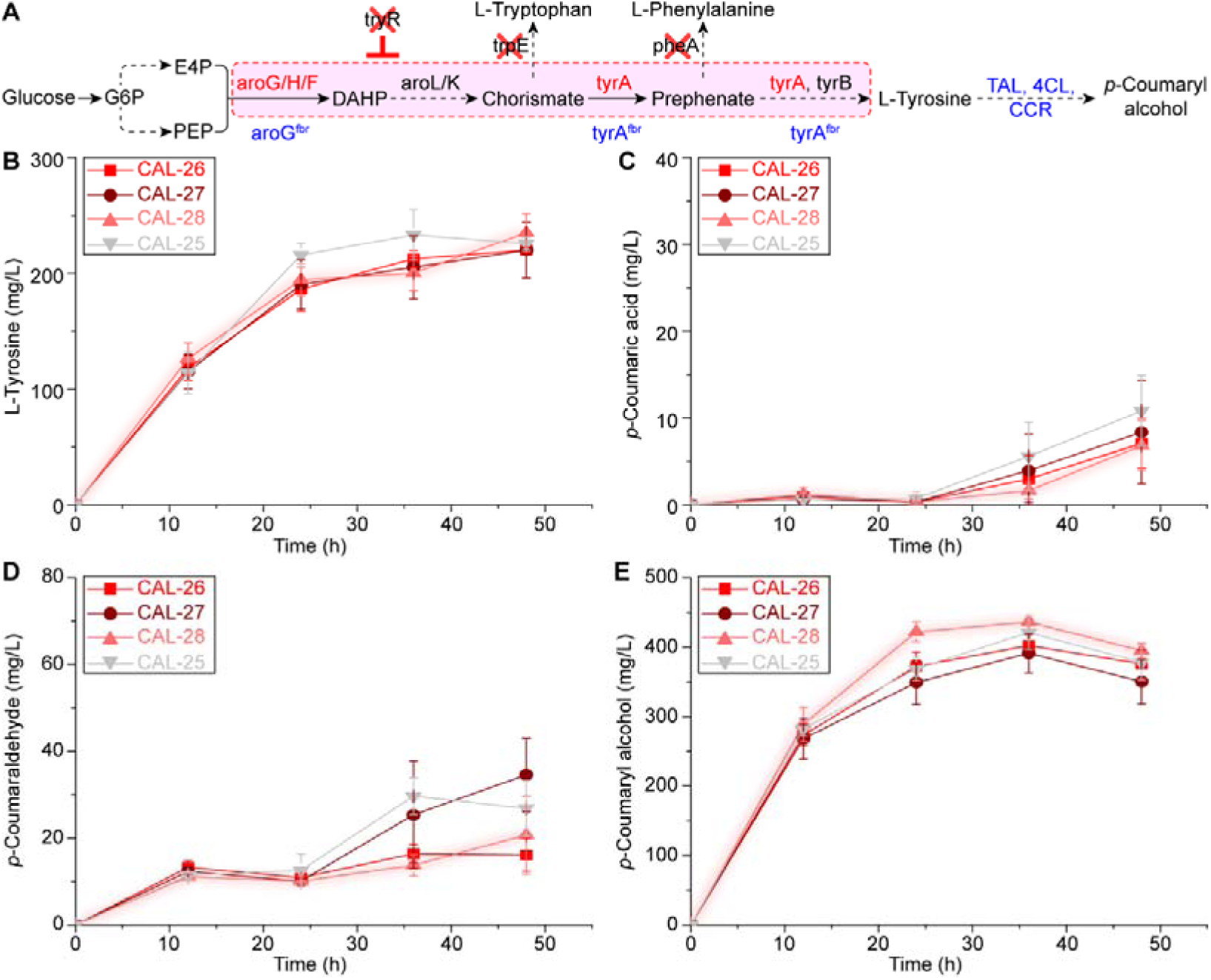
Engineering *p*-coumaryl alcohol over-accumulation by removing competing pathways. The competing pathways were blocked by deleting *pheA* and *trpE*, which lead the branched pathways to L-phenylalanine and L-tryptophan synthesis. **(A)** Schematic demonstration of the pathways. **(B)** – **(E)** The accumulations of L-tyrosine, *p*-coumaric acid, *p*-coumaraldehyde, and *p*-coumaryl alcohol.

However, this only achieved marginal improvement of *p*-coumaryl alcohol production. Strain CAL-28, containing the double deletion Δ*pheA* and Δ*trpE*, achieved 14.41% improvement at 24 h and 3.59% improvement at 36 h, compared with the parental strain CAL-25 (**Figures 4B-E**). These deletions were also tested in CAL-24, because it accumulated higher titers of *p*-coumaric acid and *p*-coumaraldehyde (**Figures 3D-G**). Similarly, only marginal improvement were obtained (**Figures S2 and S3**). The best performing strain CAL-28, which accumulated 436.28 mg/L *p*-coumaryl alcohol at 36 h, will be used in the following studies.

### 2.4 Stepwise demonstrating the synthesis of estragole and anethole

In the proposed synthesis pathway, estragole will be produced from *p*-coumaryl alcohol by the sequential activities of CFAT, EGS/APS, and OMT (**Figure 1**). When introducing CFAT into CAL-28 (**Figures 5A**), a new peak emerged in the HPLC analysis at 15.51 min (**Figure 5B**), which was deduced to be *p*-coumaryl acetate. Among the resulting strains (CAT-01 to 06, **Table S4**), CAT-02 harboring ScCFAT resulted in largest peak area (**Figure 5C**). Because *p*-coumaryl acetate standard is not commercially available, the speculated best-performing CAT-02 was used to produce chavicol. Introducing EGS/APS into CAT-02 achieved chavicol production, which was confirmed by comparing with the chavicol standard and blank control (**Figures 5D and E**). Among the resulting strains (CHA-01 to 03, **Table S4**), CHA-01 harboring MdoPhR5 produced highest chavicol titer (26.10 mg/L, **Figure 5F**). To doublecheck the activities of CFATs obtained in **Figures 5B and C**, we combined the best-performing MdoPhR5 with these CFATs, resulting in recombinant strains CHA-04 to 08 (**Table S4**). The correlation between chavicol titers and the peak areas of putative *p*-coumaryl acetate re-confirmed the previous speculation of CFAT activities (**Figures S4A** and **B**). Finally, OMT encoding genes were introduced into CHA-01, resulting in strains EST-01 to 08 (**Table S4**). Among them, EST-01, 03, and 04, which harbor PaAIMT1, ObaCVOMT1, and ObaEOMT1 respectively, achieved the production of estragole. Among them, EST-03 produced the highest estragole titer (0.64 mg/L, **Figures 5G-I**).

**Figure 5.**
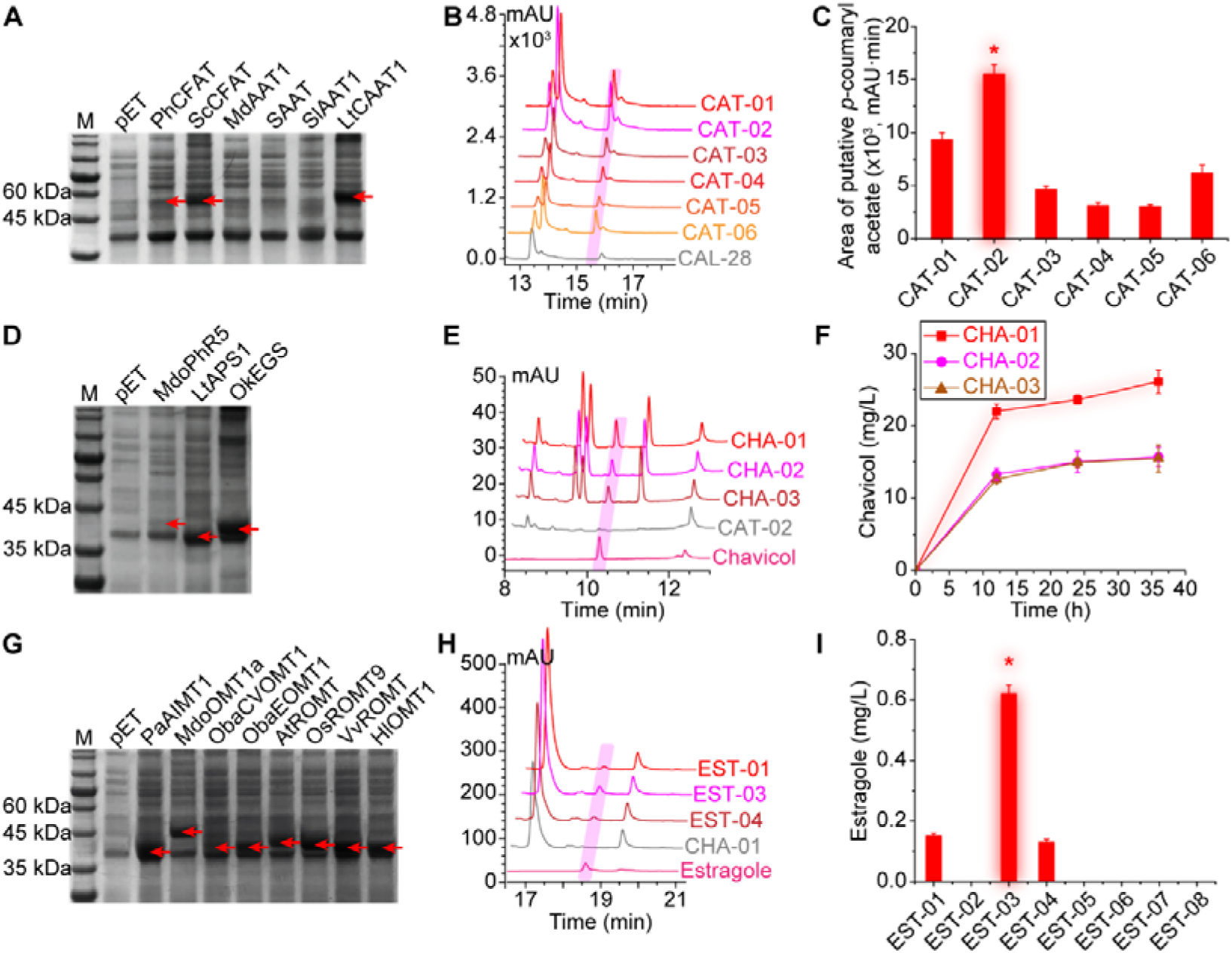
Stepwise demonstration of the estragole synthesis pathway. The estragole synthesis pathway was stepwise demonstrated on the basis of *p*-coumaryl alcohol overaccumulation. **(A)**, **(D)**, and **(G)** SDS-PAGE analysis of the overexpression of candidate CFAT, EGS/APS, and OMT. Molecular weights of recombinant proteins were as follows: PhCFAT 54.53 kDa, ScCFAT 53.74 kDa, MdAAT1 54.13 kDa, SAAT 54.11 kDa, SlAAT1 53.25 kDa, LtCAAT1 54.66 kDa, MdoPhR5 39.71 kDa, LtAPS1 37.51 kDa, OkEGS 39.12 kDa, PaAIMT1 43.04 kDa, MdoOMT1a 49.10 kDa, ObaCVOMT1 43.33 kDa, ObaEOMT1 43.65 kDa, AtROMT 43.03 kDa, OsROMT9 43.15 kDa, VvROMT 43.51 kDa, HlOMT1 42.68 kDa. Red arrows indicate the specific bands. pET, referring to *E. coli* BL21(DE3) containing empty plasmid pET28a(PB)N, was used as blank control. Other lanes were *E. coli* BL21(DE3) overexpressing corresponding genes with pET28a(PB)N. **(B)**, **(E)**, and **(H)** HPLC analysis of the production of *p*-coumaryl acetate, chavicol, and estragole. CAL-28, CAT-02, and CHA-01 were used as the blank control. **(C)** Peak area of the putative *p*-coumaryl acetate. **(F)** and **(I)** Titers of chavicol and estragole.

The anethole synthetic pathway shares CFAT with that of estragole (**Figure 1**). Screening of the candidate IGS and PPS was carried out on the basis of strain CAT-02, which accumulated highest *p*-coumaryl acetate titer. Introducing the candidate encoding genes resulted in strains ISC-01 to 03 (**Figure 6A** and **Table S4**). Compared with the parental strain, all the resulting strains produced a new peak at 10.38 min in the HPLC analysis (**Figure 6B**). As isochavicol standard is not commercially available, this new emerging peak was deduced to be isochavicol. Strain ISC-01, harboring PaAIS1, produced the largest peak area, and thus was speculated to be best-performing one (**Figure 6C**). The production of anethole was carried out on the basis of ISC-01 by introducing the OMT encoding genes (**Figure 5G**). Among the resulting strains (ANT-01 to 08, **Table S4**), only ANT-01 and ANT-04, containing PaAIMT1 and ObaEOMT1 respectively, produced anethole. The production was confirmed by comparing with the anethole standard and blank control (**Figure 6D**). Among them, ANT-04 produced higher anethole titer (2.46 mg/L) (**Figures 6E**). To doublecheck the performance of the candidate IGS and PPS, we combined LtPPS1 and PhIGS1 with OMTs, resulting in strains ANT-09 to 24 (**Table S4**). Similar to those containing PaAIS1 (ANT-01 to 08, **Figures 6E**), only the combinations with PaAIMT1 and ObaEOMT1 (ANT-09, ANT-12, ANT-17, and ANT-20) produced anethole (**Figures S5A and B**). Moreover, the correlation between anethole titers and the peak areas of deduced isochavicol re-confirmed the previous speculation of the IGS and PPS activities (**Figures S5C**).

**Figure 6.**
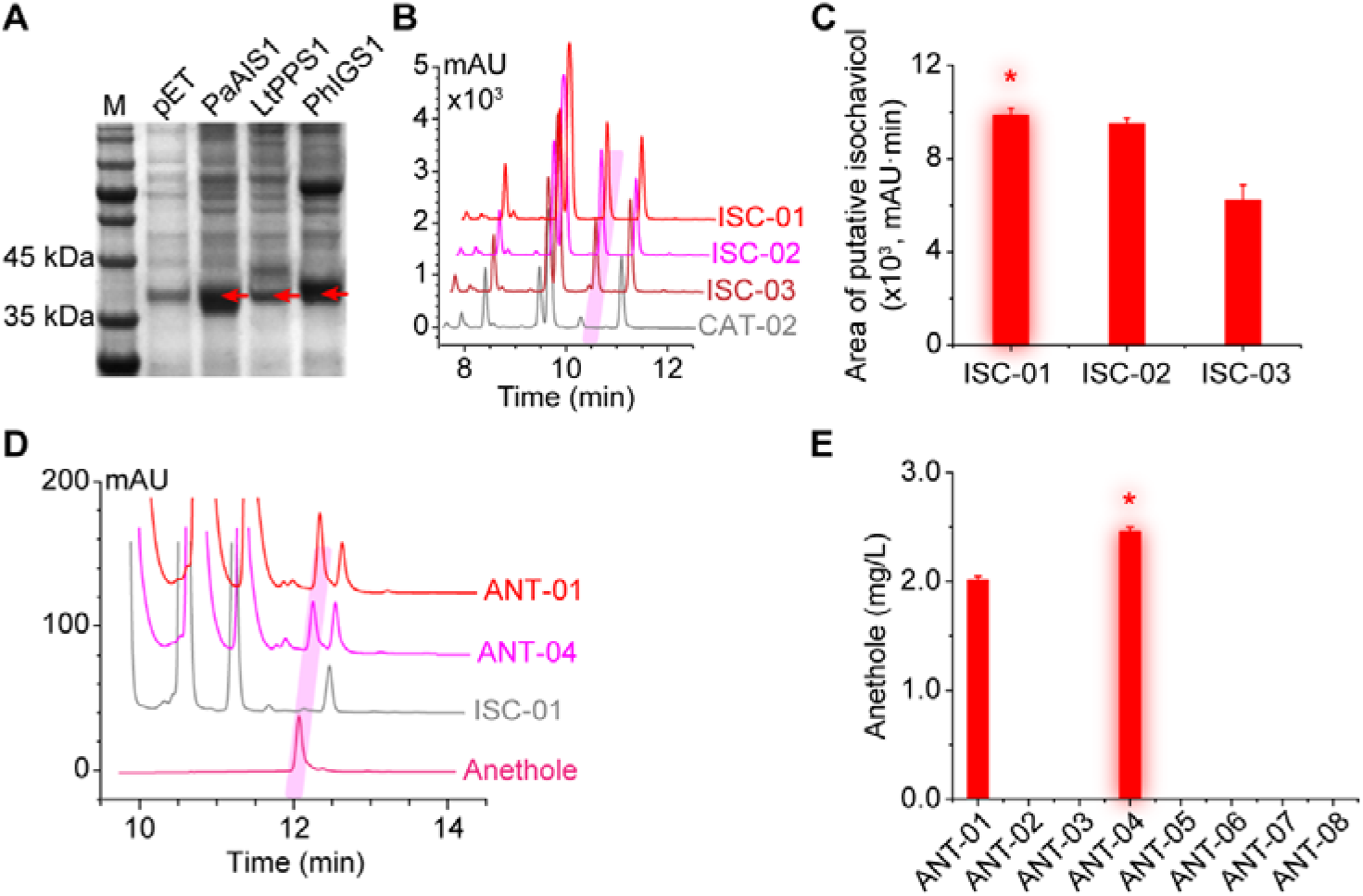
Stepwise demonstration of the anethole synthesis pathway. The anethole synthesis pathway was stepwise demonstrated on the basis of *p*-coumaryl acetate production. **(A)** SDS-PAGE analysis of the overexpression of candidate IGS and PPS. Molecular weights of recombinant proteins were as follows: PaAIS1 39.60 kDa, LtPPS1 39.19 kDa, PhIGS1 39.41 kDa. Red arrows indicate the specific bands. pET, referring to *E. coli* BL21(DE3) containing empty plasmid pET28a(PB)N, was used as blank control. Other lanes were *E. coli* BL21(DE3) overexpressing corresponding genes with pET28a(PB)N. **(B)** HPLC analysis of the isochavicol production. CAT-02 was used as blank control. **(C)** Peak areas of the putative isochavicol. **(D)** HPLC analysis of the anethole production. ISC-01 was used as the blank control. **(E)** Anethole titers produced by ANT-01 to 08.

### 2.5 Improving the production of estragole and anethole by pathway engineering and improving SAM availability

Based on the achieving of *de novo* biosynthesis, we tried to improve the production of estragole and anethole by debottlenecking the synthesis pathway. Stepwise increasing the gene copy numbers, resulting in strains EST-09 to 16 and ANT-25 to 32 (**Table S4**), indicated that methylation serves as the rate limiting node of both pathways. Increasing the copy number of ObaCVOMT1 and ObaEOMT1 to 3 improved the titers to 0.73 mg/L (estragole, **Figure 7A**) and 3.67 mg/L (anethole, **Figure 7B**), respectively. The dramatic decrease of estragole titer (0.62 mg/L) compared with its precursor chavicol (26.09 mg/L) also indicated that the methylation serves as the limiting node (**Figures 5F and I**).

**Figure 7.**
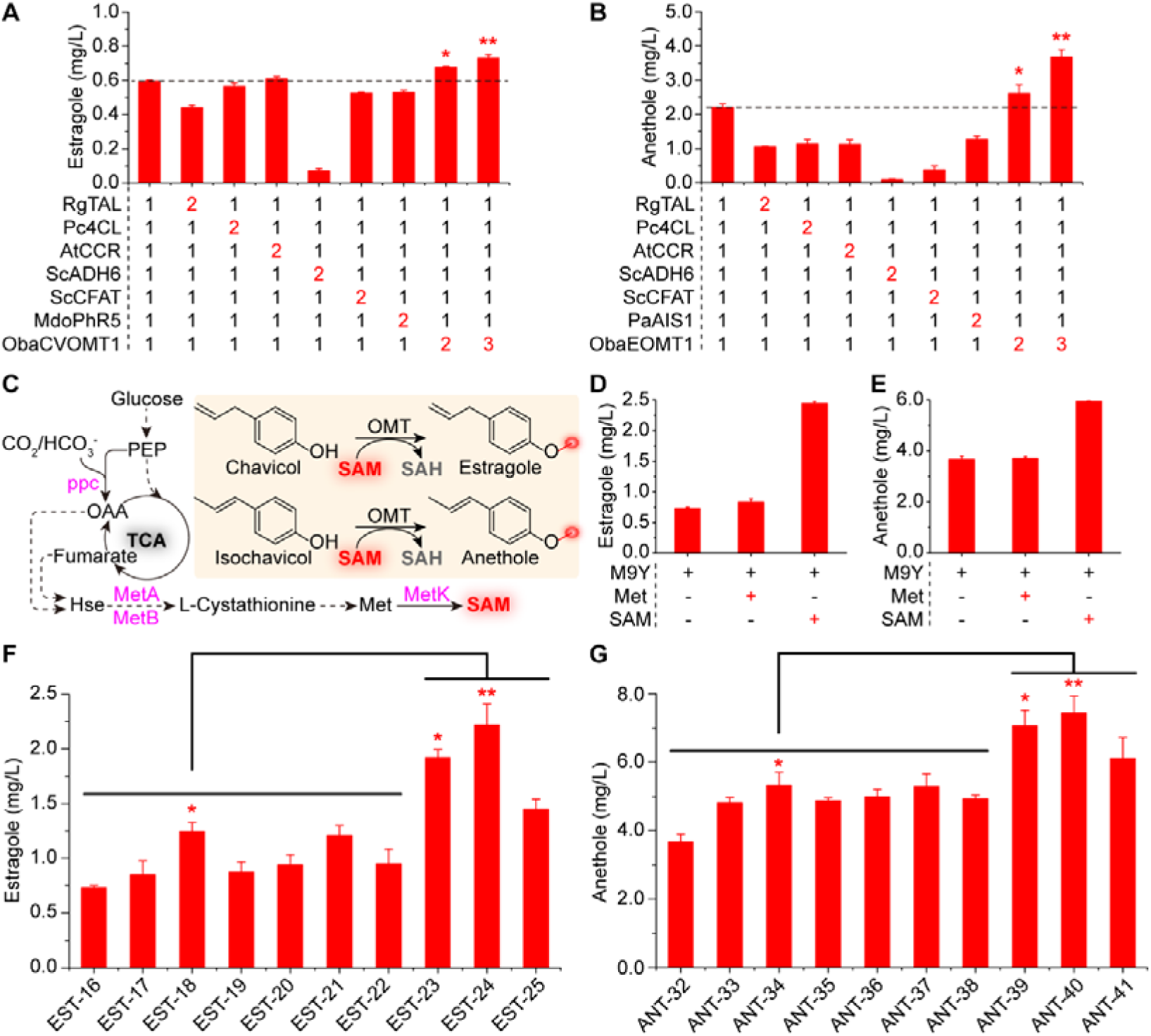
Improving the production by pathway engineering and improving SAM availability Debottlenecking the synthesis pathway of estragole. **(A)** and anethole **(B)** by stepwise increasing gene copy number. The numbers along x axis indicate the gene copy numbers. **(C)** The synthesis pathway of SAM and the synthesis of estragole and anethole using SAM as methyl group donor. Magenta indicates the potential limiting node. Improving the production of estragole **(D)** and anethole **(E)** by feeding L-methionine and SAM. Improving the production of estragole **(F)** and anethole **(G)** by enhancing SAM synthesis.

SAM serves as the direct methyl group donor of the methylation (**Figure 7C**). However, because of the strict homeostasis regulation, its low intracellular availability hampers the over-production of SAM-dependent methylated products by the MCF ^8^. This was confirmed by the heterogenous feeding of SAM and its precursor L-methionine. Feeding 2 mM SAM improved the titers of estragole and anethole by 234.25% and 62.13%, respectively (**Figures 7D and E**). *S*-Adenosylmethionine synthetase catalyzes the final step of SAM synthesis, where L-methionine and ATP are converted into SAM, releasing pyrophosphate and phosphate. This step has been proven the key node to enhance the over-production of SAM in *E. coli* ^25^. We introduced 6 *S*-adenosylmethionine synthetase encoding genes from different organisms (**Table S1**) into the best-performing EST-16 and ANT-32, respectively. All the resulting strains produced higher titers of estragole (EST-17 to 22, **Table S4**) and anethole (ANT-33 to 38, **Table S4**) than their parental strains (**Figures 7F and G**). Among them, EST-18 and ANT-34, both of which contain *S*-adenosylmethionine synthetase encoding gene *BcMetK* from *Bacillus cereus*, produced the highest titers of estragole (1.25 mg/L) and anethole (5.31 mg/L), respectively. In an other study, we discovered that the overexpression of *ppc* (encoding phosphoenolpyruvate carboxylase), *metA* (encoding bifunctional homoserine *O*-succinyltransferase/*O*-acetyltransferase), and *metB* (encoding cystathionine γ-synthase) was beneficial for the production of SAM-dependent products. Consequently, we overexpressed these genes in EST-18 and ANT-34, respectively. The resulting strains produced higher titers of estragole (EST-23 to 25, **Table S4**) and anethole (ANT-39 to 41, **Table S4**) than their parental strains (**Figures 7F and G**). Among these strains, EST-24 and ANT-40, both of which co-overexpressed *BcMetK* and *metA*, produced the highest titers of estragole (2.22 mg/L) and anethole (7.44 mg/L). These represent improvements of 77.60% and 40.11% than their parental strains, and 246.88% and 202.44% than the initial strains.

Estragole and anethole are among the major components of the essential oils extracted from basil (*Ocimum basilicum* L.), anise (*Pimpinella anisum*), star anise (*Illicium verum*), and fennel (*Foeniculum vulgare*), and have been substantially used in the industries of food, cosmetics, pharmaceutics, and agriculture ^6, 26^. Most commercial estragole and anethole are obtained by extracting from their original plants ^27^. In instability of quality and resource hinders their applications ^3^. In this study, we stepwise demonstrated the synthetic pathway and achieved the *de novo* synthesis in microbial chassis for the first time. The methylation was determined to be the rate limiting step through the synthetic pathway. The productions were enhanced by 246.88% (estragole) and 202.44% (anethole) by pathway debottlenecking and enhancing cofactor SAM availability. Intriguingly, despite the relative low titers, the scents of estragole and anethole were pretty strong as long as they were produced. When the complete pathways were assembled, the scents were smelt nearby the incubator during the shaking flask fermentation. This study represents the first *de novo* biosynthesis of estragole and anethole in microbial chassis, and the first demonstration of microbial synthesis of propenyl phenols. Moreover, it provided a sustainable alternative approach to obtain the natural flavoring and aroma chemicals.

## 3. Materials and methods

### 3.1 Genes, plasmids, and strains

*Escherichia coli* JM109 was used for gene cloning, plasmid construction, and plasmid propagation. *E. coli* BL21(DE3) was used for gene expression, pathway analysis, and the production of estragole, anethole and their intermediates. The ePathBricks plasmids pACM4, pCDM4, and pET28a(PB)N were developed in our previous studies ^14, 17^, and used for gene expression and pathway construction. Plasmids used and developed in this study were listed in Supplementary Information **Table S3**. The genes used in this study were listed in **Table S1**. The strains used and developed in this study were listed in **Table S4**.

### 3.2 Cell culture and fermentation

LB medium was used for molecular biology manipulations. M9Y medium was used for protein expression analysis and the production of estragole, anethole, and their intermediates. The fermentation was carried out in 250 mL shaking flask containing 25 mL M9Y medium. M9Y medium contains 98% (v/v) M9Y minimal, 1% (v/v) 100× salt I, and 1% (v/v) 100× salt II. M9Y minimal contains 5 g/L glucose, 10 g/L glycerol, 5 g/L yeast extract, 0.5 g/L NaCl, 17.1 g/L Na_2_HPO_4_·12H_2_O, 3 g/L KH_2_PO_4_, 2 g/L NH_4_Cl. For the culture of phenylalanine and tryptophan auxotrophic strains, 100 mg/L L-phenylalanine and 40 mg/L L-tryptophan were added. The 100× salt I contains 0.1 M MgSO_4_·7H_2_O, 1 mM FeSO_4_·7H_2_O, and the 100× salt II contains 0.01 M CaCl_2_·2H_2_O. M9Y minimal, 100× salt I, and 100× salt II were autoclaved separately at 115°C for 15 min, and mixed before use ^28^. Antibiotic(s) was added as necessary ^14, 17^. Final concentration of 2 mM L-methionine and SAM were added for the SAM availability assays. Biological reagent grade chemicals were purchased from Sangon Biotech (Shanghai) Co., Ltd. (Shanghai, China).

### 3.3 Molecular biology

The genes from *Saccharomyces cerevisiae*, *Yarrowia lipolytica*, and *Escherichia coli* were amplified from the corresponding genome DNA, while other genes were codon-optimized and synthesized by Sangon Biotech (Shanghai) Co., Ltd. (Shanghai, China). The primers were listed in Supplementary Information **Table S2**. For gene overexpression, the genes were subcloned into pET28a(PB)N between *Bam*HI and *Xho*I sites, pACM4 between *Eco*RV and *Xho*I sites, or pCDM4 between *Nde*I and *Xho*I sites. The subcloning of genes synthesized by Sangon Biotech was carried out by restriction digestion and T4 ligation, while the subcloning of other genes was carried out by using the One-step cloning kit (Vazyme Biotech, Nanjing, China). The gene co-overexpression and pathway construction were carried out following the ePathBricks method. Specifically, the donor fragment was digested with *Avr*II and *Sal*I enzymes, and the acceptor plasmid was digested with *Nhe*I and *Sal*I enzymes. The pathways were assembled into monocistronic form by the subsequent T4 ligation^17^.

The gene knockout was carried out using the CRISPR-Cas9 system developed by Jiang *et al* ^29^. Plasmids pCas and pTargetF were kindly donated by Dr. Yang. Specifically, the N20 sequence of pTargetF was changed by amplifying the entire plasmid and circularization using the One-step cloning kit. The deletion fragment containing up- and down-stream homologous arms were PCR amplified from the genome DNA of *E. coli* BL21(DE3), fused together using overlapping PCR, and subcloned into pUC57 vector between *Bam*HI and *Hind*III sites. The deletion fragments were checked by Sanger sequencing and amplified from the vector for electroporation. The manipulations were carried out following the guidance ^29^. Gene knockout was confirmed using the diagnostic PCR analysis. Primers used in this part were in **Table S2**. The plasmids used and developed in this part were in **Table S3**.

Protein overexpression was confirmed by using the SDS-PAGE analysis, and compared with the standard ladder and blank control. It should be noted that the N-terminal His-tag made the recombinant protein larger than the natural one.

### 3.4 Analytical method

Samples were mixed with equal volume of ethanol, and shook in 1.5-mL Eppendorf tubes with 0.1-mm diameter glass beads to break cell wall. After centrifugation and filtration, samples were injected into HPLC and LC/MS for analysis.

HPLC analysis was carried out with an Agilent 1260 system. Samples were separated with an Eclipse Plus C18 (4.6 mm×250 mm×5 μm) column, and detected with a VWD detector (λ=286 nm). The oven temperature was 40°C. Flow rate was 1 mL/min. Inject volume was 5 μL. The mobile phase A was water with 0.1% (v/v) acetate, and mobile phase B was methanol with 0.1% (v/v) acetate. The gradient (B%) was as follows: 0 min 0%, 5 min 65%, 7 min 80%, 9 min 100%, 12 min 100%, 14 min 50%, 18 min 0%, maintaining 0% until 20 min.

The LC/MS analysis was carried out with an Agilent 6430 Triple Quadrupole LC/MS System. Samples were separated with an Eclipse Plus C18 (4.6 mm×250 mm×5 μm) column, and detected with a DAD detector (λ=286 nm). The oven temperature was 25°C. Flow rate was 0.7 mL/min. Inject volume was 5 μL. The mobile phase A was water with 0.1% (v/v) acetate, and mobile phase B was methanol with 0.1% (v/v) acetate. The gradient (B%) was as follows: 0 min 0%, 8 min 65%, 11 min 80%, 15 min 100%, 20 min 100%, 22 min 50%, 28 min 0%, maintaining 0% until 30 min. Electrospray ionization (ESI) ion source and negative mode were used^30^.

Analytical standards were purchased from Shanghai Yuanye Bio-Technology Co., Ltd.

## Author contributions

YL conceived the topic. YL designed the experiments and analyzed the data. JC, YL, and SD performed the experiments. YL and PX drafted and revised the manuscript. SG, YX, AZ, SW, WW, YX, and JX gave suggestions.

## Ethical approval

This article does not contain any studies with human participants or animals performed by any of the authors.

## Supporting information

The Supplementary Information file includes supporting information as follows: Figure S1 Analysis of the expression of CCR and CAD and the production of *p*-coumaryl alcohol; Figure S2 Diagnostic PCR analysis of gene knockout; Figure S3 Improving *p*-coumaryl alcohol accumulation by blocking competing pathways in CAL-24; Figure S4 Rechecking the activities of coniferyl alcohol acyltransferase; Figure S5 Rechecking the activities of PPS and IGS; Table S1 Genes used in this study; Table S2 Primers used in this study; Table S3 Plasmids used and developed in this study; Table S4 Strains used and developed in this study.

