## Supplementary Information for "Demonstrating and engineering the *de novo* synthesis of two aromatic chemicals estragole and anethole in *Escherichia coli*"

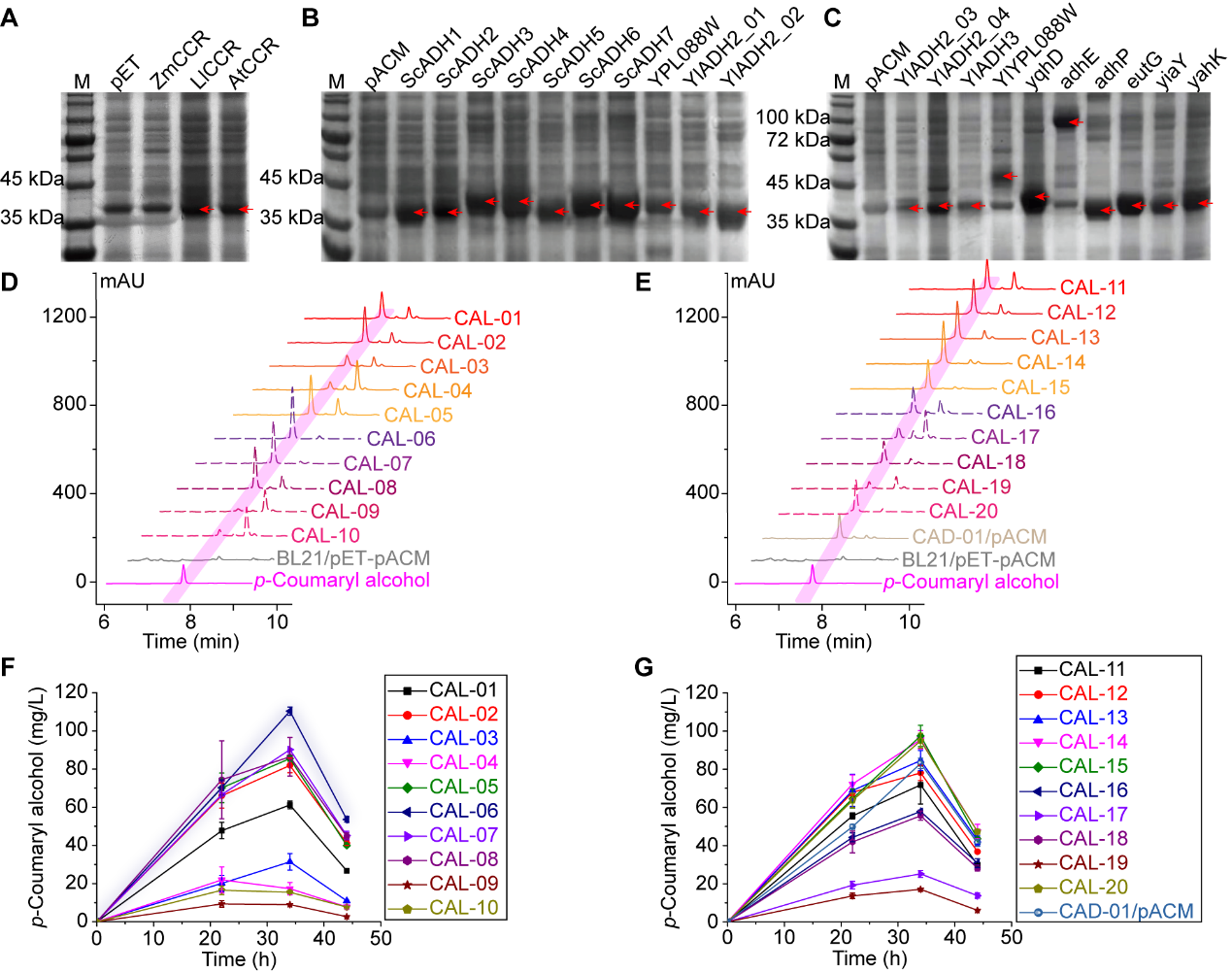

Figure S1 Analysis of the expression of CCR and CAD and the production of *p*-coumaryl alcohol.

**(A)** SDS-PAGE analysis of the CCR expression. Calculated molecular weights are as follows: ZmCCR 38.00 kDa, LlCCR 36.52 kDa, AtCCR 36.61 kDa. pET refers to *E. coli* BL21(DE3) containing empty plasmid pET28a(PB)N, and was used as the blank control. Other lanes are *E. coli* BL21(DE3) overexpressing corresponding genes with pET28a(PB)N. **(B)** and **(C)** SDS-PAGE analysis of the expression of alcohol dehydrogenases and aldehyde reductase. Calculated molecular weights are as follows: ScADH1 36.71 kDa, ScADH2 36.60 kDa, ScADH3 40.23 kDa, ScADH4 41.01 kDa, ScADH5 37.51 kDa, ScADH6 39.48 kDa, ScADH7 39.21 kDa, YPL088W 39.55 kDa, YlADH2_01 36.92 kDa, YlADH2_02 36.85 kDa, YlADH2_03 35.25 kDa, YlADH2_04 37.18 kDa, YlADH3 37.82 kDa, YlYPL088W 48.09 kDa, yqhD 41.96 kDa, adhE 95.99 kDa, adhP 35.24 kDa, eutG 40.90 kDa, yiaY 40.22 kDa, yahK 37.84 kDa. pACM refers to *E. coli* BL21(DE3) containing empty plasmid pACM4, and was used as the blank control. Other lanes are *E. coli* BL21(DE3) overexpressing corresponding genes with pACM4. Red arrows indicate the recombinant enzymes. **(D)** and **(E)** HPLC analysis of *p*-coumaryl alcohol production by strains containing different alcohol dehydrogenases. **(F)** and **(G)** The *p*-coumaryl alcohol titers produced by strains containing different alcohol dehydrogenases.

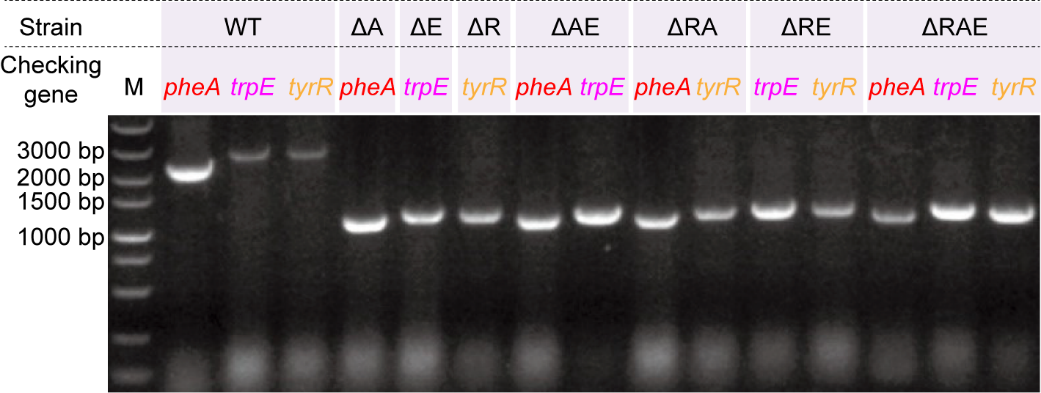

Figure S2 Diagnostic PCR analysis of gene knockout.

The gene knockout was validated by using diagnostic PCR analysis. The specific bands of wild type *pheA*, *trpE*, and *tyrR* were 2359-bp, 2760-bp, and 2714-bp. The specific bands of the mutant *pheA*, *trpE*, and *tyrR* were 1198-bp, 1197-bp, and 1172-bp. WT refers to the wild type *E. coli* BL21(DE3).

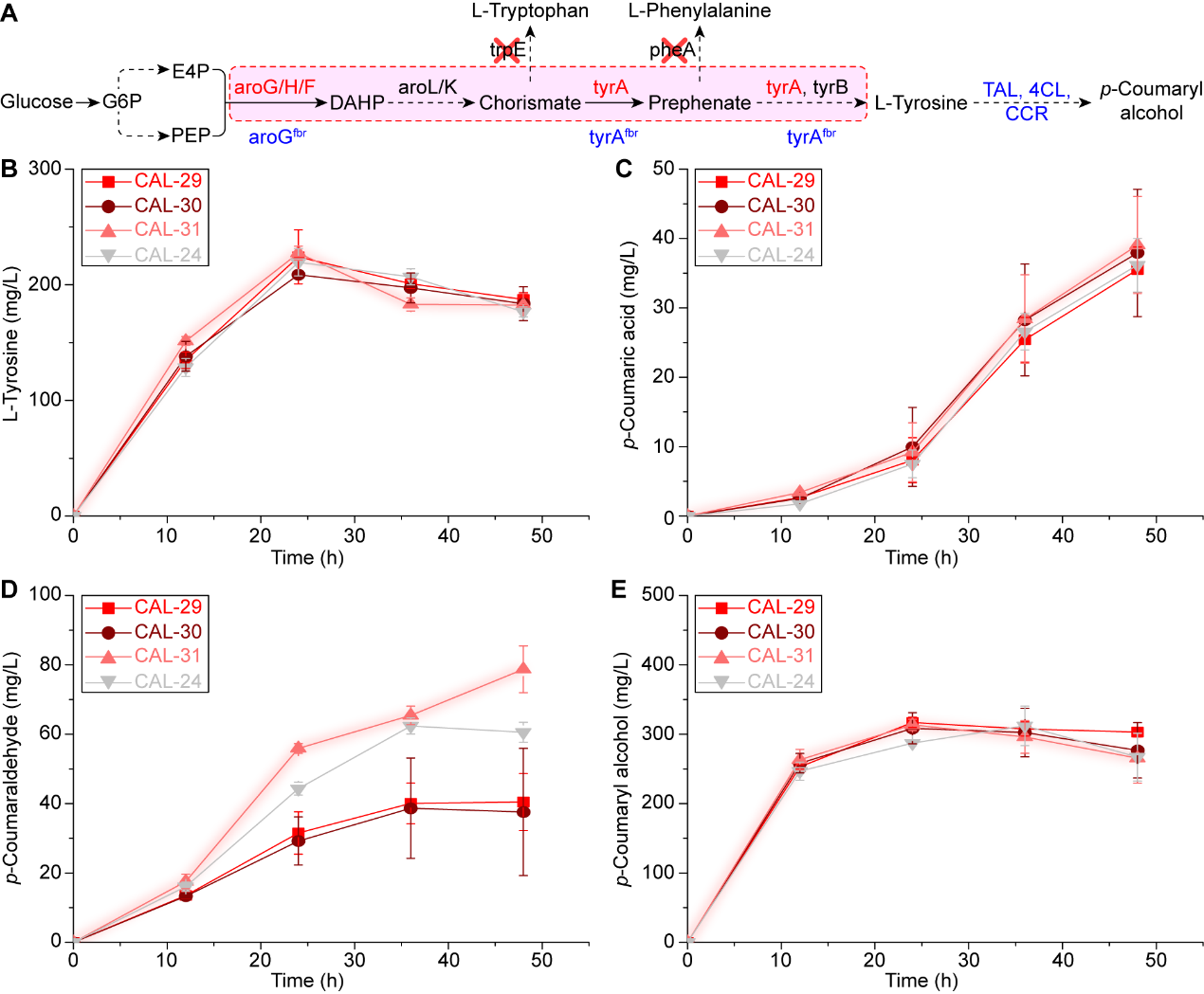

Figure S3 Improving *p*-coumaryl alcohol accumulation by blocking competing pathways in CAL-24.

The competing pathways were blocked by deleting *pheA* and *trpE*, which lead the branched pathways to L-phenylalanine and L-tryptophan synthesis. **(A)** Schematic demonstration of the pathways. **(B)** – **(E)** The accumulations of L-tyrosine, *p*-coumaric acid, *p*-coumaraldehyde, and *p*-coumaryl alcohol.

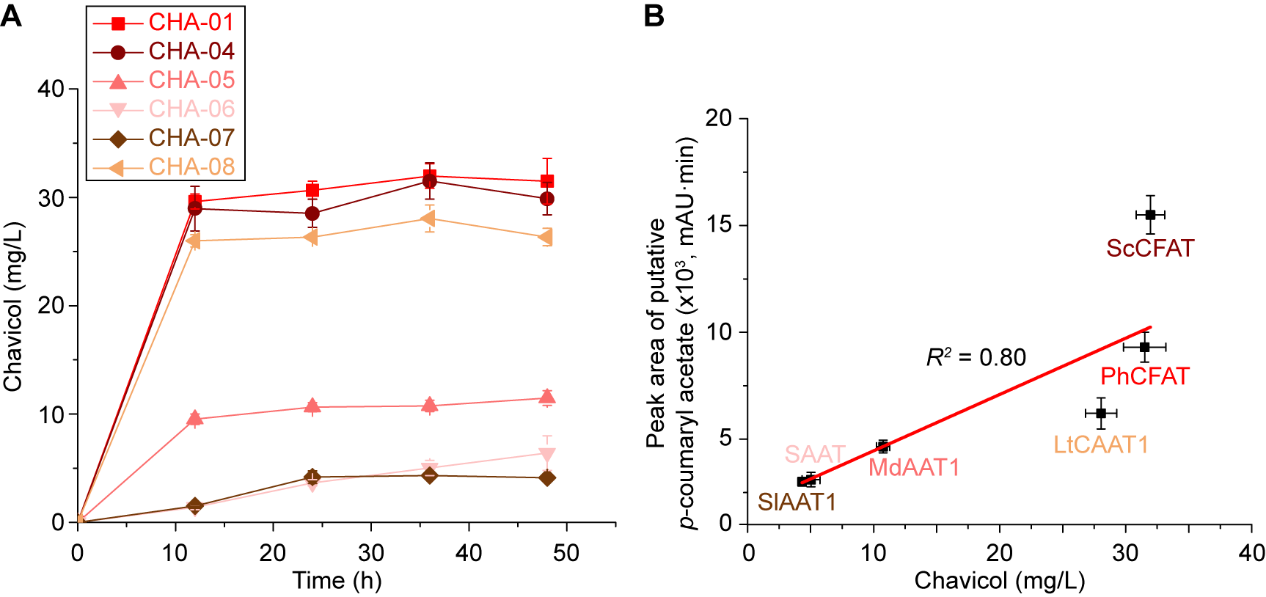

Figure S4 Rechecking the activities of coniferyl alcohol acyltransferase.

The activities of coniferyl alcohol acyltransferase (CFAT) were rechecked by measuring the titers of chavicol. **(A)** The chavicol titers produced by strains expressing different CFATs. **(B)** The correlation between chavicol titers and the peak areas of the putative *p*-coumaryl acetate.

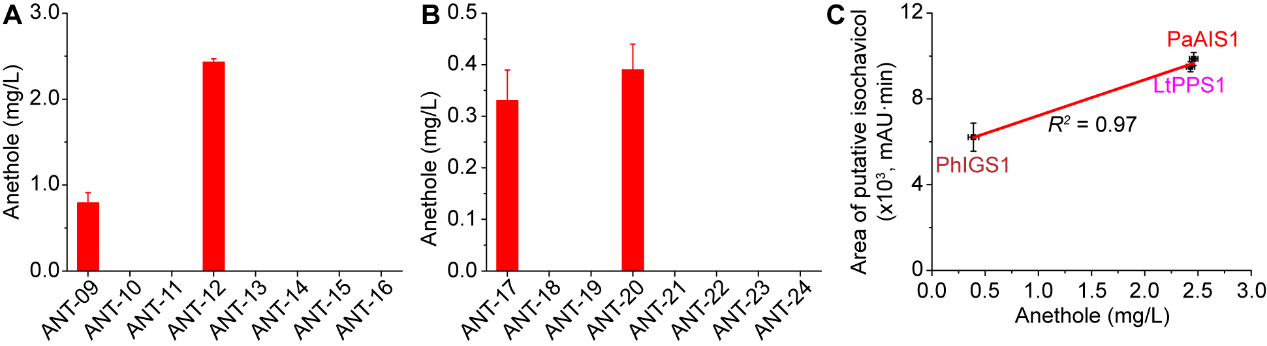

Figure S5 Rechecking the activities of PPS and IGS.

The activities of propenyl phenol synthase (PPS) and isoeugenol/isochavicol synthase (IGS) were rechecked by measuring the titers of anethole. **(A)** The anethole titers produced by strains expressing LtPPS1 and different OMTs. **(B)** The anethole titers produced by strains expressing PhIGS1 and different OMTs. **(C)** The correlation between anethole titers and the peak areas of the putative isochavicol.

Table S1 Genes used in this study

| **#** | **Gene** | **Encoding enzyme** | **Source organism** | **Database access No.** | **Reference** |
| --- | --- | --- | --- | --- | --- |
| 1 | *RgTAL* | Tyrosine ammonia-lyase | *Rhodotorula toruloides* | NCBI: AAA33883 | ^1^ |
| 2 | *Pc4CL* | *p*-Coumarate-CoA ligase | *Petroselinum crispum* (parsley) | UniProtKB: P14912 | ^1^ |
| 3 | *AtCCR* | Cinnamoyl-CoA reductase | *Arabidopsis thaliana* | NCBI: AEE36454.1 | ^2^ |
| 4 | *ZmCCR* | Cinnamoyl-CoA reductase | *Zea mays* | NCBI: O82726 | ^3^ |
| 5 | *LlCCR* | Cinnamoyl-CoA reductase | *Leucaena leucocephala* | NCBI: CAK22319.1 | ^4, 5^ |
| 6 | *ScADH1* | Alcohol dehydrogenase | *Saccharomyces cerevisiae* | NCBI: 854068 | ^6^ |
| 7 | *ScADH2* | Alcohol dehydrogenase | *Saccharomyces cerevisiae* | NCBI: 855349 | ^7^ |
| 8 | *ScADH3* | Alcohol dehydrogenase | *Saccharomyces cerevisiae* | NCBI: 855107 | ^8^ |
| 9 | *ScADH4* | Alcohol dehydrogenase | *Saccharomyces cerevisiae* | NCBI: 852636 | ^8^ |
| 10 | *ScADH5* | Alcohol dehydrogenase | *Saccharomyces cerevisiae* | NCBI: 852442 | ^8^ |
| 11 | *ScADH6* | Alcohol dehydrogenase | *Saccharomyces cerevisiae* | NCBI: 855368 | ^4, 9^ |
| 12 | *ScADH7* | Alcohol dehydrogenase | *Saccharomyces cerevisiae* | NCBI: 850469 | ^8^ |
| 13 | *YPL088W* | Aldo-keto reductase | *Saccharomyces cerevisiae* | NCBI: 856017 |  |
| 14 | *YlADH2_01* | Alcohol dehydrogenase | *Yarrowia lipolytica* | NCBI: YALI0A16379g |  |
| 15 | *YlADH2_02* | Alcohol dehydrogenase | *Yarrowia lipolytica* | NCBI: YALI0D25630g |  |
| 16 | *YlADH2_03* | Alcohol dehydrogenase | *Yarrowia lipolytica* | NCBI: YALI0E17787g |  |
| 17 | *YlADH2_04* | Alcohol dehydrogenase | *Yarrowia lipolytica* | NCBI: YALI0A15147g |  |
| 18 | *YlADH3* | Alcohol dehydrogenase | *Yarrowia lipolytica* | NCBI: YALI0E07766g |  |
| 19 | *YlYPL088W* | Aldo-keto reductase | *Yarrowia lipolytica* | NCBI: YALI0C12595g |  |
| 20 | *yqhD* | NADPH-dependent aldehyde reductase | *Escherichia coli* | NCBI: 947493 | ^10^ |
| 21 | *adhE* | Alcohol dehydrogenase | *Escherichia coli* | NCBI: 945837 |  |
| 22 | *adhP* | Alcohol dehydrogenase | *Escherichia coli* | NCBI: 946036 | ^11^ |
| 23 | *eutG* | Alcohol dehydrogenase | *Escherichia coli* | NCBI: 946233 | ^12^ |
| 24 | *yiaY* | Alcohol dehydrogenase | *Escherichia coli* | NCBI: 948102 | ^12^ |
| 25 | *yahK* | Alcohol dehydrogenase | *Escherichia coli* | NCBI: 944975 | ^13^ |
| 26 | *PhCFAT* | Coniferyl alcohol acyltransferase | *Petunia × hybrida* | DQ767969 | ^14^ |
| 27 | *ScCFAT* | Coniferyl alcohol acyltransferase | *Schisandra chinensis* | OM804184 | ^15^ |
| 28 | *MdAAT1* | Alcohol acyl transferase | *Malus × domestica* | KC291129 | ^16^ |
| 29 | *SAAT* | Alcohol acyl transferase | *Fragaria × ananassa* | AAG13130 | ^16^ |
| 30 | *SlAAT1* | Alcohol acyl transferase | *Solanum lycopersicum* | KM975322 | ^16^ |
| 31 | *LtCAAT1* | Alcohol acyl transferase | *Larrea tridentata* | KF543260 | ^17^ |
| 32 | *MdoPhR5* | Eugenol/chavicol synthase | *Malus domestica* | KX577779 | ^16^ |
| 33 | *LtAPS1* | Eugenol/chavicol synthase | *Larrea tridentata* | KF543262 | ^17^ |
| 34 | *OkEGS* | Eugenol/chavicol synthase | *Ocimum kilimandscharicum* | AOC97443.1 | ^18^ |
| 35 | *PhIGS1* | Isoeugenol/isochavicol synthase | *Petunia × hybrida* | Q15GI3.1 |  |
| 36 | *LtPPS1* | Isoeugenol/isochavicol synthase | *Larrea tridentata* | KF543264 | ^17^ |
| 37 | *PaAIS1* | Isoeugenol/isochavicol synthase | *Pimpinella anisum* | EU925388 | ^19^ |
| 38 | *PaAIMT1* | *O*-Methyltransferase | *Pimpinella anisum* | EU925389 | ^19^ |
| 39 | *MdoOMT1a* | *O*-Methyltransferase | *Malus domestica* | KM516782 | ^16, 20^ |
| 40 | *ObaCVOMT1* | *O*-Methyltransferase | *Ocimum basilicum* | AF435007 | ^21^ |
| 41 | *ObaEOMT1* | *O*-Methyltransferase | *Ocimum basilicum* | AF435008 | ^21^ |
| 42 | *AtROMT* | *O*-Methyltransferase | *Arabidopsis thaliana* | UniProtKB: Q9FK25 | ^22^ |
| 43 | *OsROMT9* | *O*-Methyltransferase | *Oryza sativa* subsp. japonica (Rice) | UniProtKB: Q6ZD89 |  |
| 44 | *VvROMT* | *O*-Methyltransferase | *Vitis vinifera* | UniProtKB: B6VJS4 | ^23^ |
| 45 | *HlOMT1* | *O*-Methyltransferase | *Humulus lupulus* (European hop) | UniProtKB: B0ZB55 | ^24^ |
| 46 | *tyrA*^fbr^ | Feedback-inhibition resistant chorismate mutase/prephenate dehydrogenase, containing M53I and A354V mutations | *Escherichia coli* | NCBI: 947115 | ^25^ |
| 47 | *aroG*^fbr^ | Feedback-inhibition resistant 3-deoxy-D-arabino-heptulosonate-7-phosphate synthase, containing a D146N mutation | *Escherichia coli* | NCBI: 945605 | ^26^ |
| 48 | EcMetK | *S*-Adenosylmethionine synthetase | *Escherichia coli* | KEGG: b2942 |  |
| 49 | BcMetK | *S*-Adenosylmethionine synthetase | *Bacillus cereus* | KEGG: BC4761 |  |
| 50 | SaMetK | *S*-Adenosylmethionine synthetase | *Staphylococcus aureus* | KEGG: SA1608 |  |
| 51 | BsMetK | *S*-Adenosylmethionine synthetase | *Bacillus subtilis* | KEGG: BSU30550 |  |
| 52 | ScSAM1 | *S*-Adenosylmethionine synthetase | *Saccharomyces cerevisiae* | KEGG: YLR180W |  |
| 53 | ScSAM2 | *S*-Adenosylmethionine synthetase | *Saccharomyces cerevisiae* | KEGG: YDR502C |  |
| 54 | ppc | Phosphoenolpyruvate carboxylase | *Escherichia coli* | KEGG: b3956 |  |
| 55 | metA | Bifunctional homoserine *O*-succinyltransferase/*O*-acetyltransferase | *Escherichia coli* | KEGG: b4013 |  |
| 56 | metB | Cystathionine γ-synthase | *Escherichia coli* | KEGG: b3939 |  |

Table S2 Primers used in this study

| **#** | **Primer** | **Sequence (5’-3’)** | **Annotation** |
| --- | --- | --- | --- |
| 1 | ScADH1-pACM F | atggcagatctcaattggtctatcccagaaactcaaaaagg | Constructing plasmid pACM4-ScADH1 |
| 2 | ScADH1-pACM R | AGCGGTTTCTTTACCAGATTATTTAGAAGTGTCAACAACG |  |
| 3 | ScADH2-pACM F | atggcagatctcaattggtctattccagaaactcaaaaagc | Constructing plasmid pACM4-ScADH2 |
| 4 | ScADH2-pACM R | AGCGGTTTCTTTACCAGATTATTTAGAAGTGTCAACAACG |  |
| 5 | ScADH3-pACM F | atggcagatctcaattggttgagaacgtcaacattgttcac | Constructing plasmid pACM4-ScADH3 |
| 6 | ScADH3-pACM R | AGCGGTTTCTTTACCAGATTATTTACTAGTATCGACGACG |  |
| 7 | ScADH4-pACM F | atggcagatctcaattggtcttccgttactgggttttac | Constructing plasmid pACM4-ScADH4 |
| 8 | ScADH4-pACM R | AGCGGTTTCTTTACCAGATTAATATTCATAGGCTTTCTTG |  |
| 9 | ScADH5-pACM F | atggcagatctcaattggccttcgcaagtcattcctgaaaaac | Constructing plasmid pACM4-ScADH5 |
| 10 | ScADH5-pACM R | AGCGGTTTCTTTACCAGATCATTTAGAAGTCTCAACAAC |  |
| 11 | ScADH6-pACM F | atggcagatctcaattggtcttatcctgagaaatttgaag | Constructing plasmid pACM4-ScADH6 |
| 12 | ScADH6-pACM R | AGCGGTTTCTTTACCAGACTAGTCTGAAAATTCTTTGTCG |  |
| 13 | ScADH7-pACM F | atggcagatctcaattggctttacccagaaaaatttcagg | Constructing plasmid pACM4-ScADH7 |
| 14 | ScADH7-pACM R | AGCGGTTTCTTTACCAGACTATTTATGGAATTTCTTATC |  |
| 15 | YPL088W-pACM F | atggcagatctcaattgggttttagttaagcaggtaagac | Constructing plasmid pACM4-YPL088W |
| 16 | YPL088W-pACM R | AGCGGTTTCTTTACCAGATTAACATCTTTGCCTCTGGGG |  |
| 17 | YlADH2_01-pACM F | atggcagatctcaattggaccaccatccccaagacccag | Constructing plasmid pACM4-YlADH2_01 |
| 18 | YlADH2_01-pACM R | AGCGGTTTCTTTACCAGATTACTTGGAGCAGTCCAGAACG |  |
| 19 | YlADH2_02-pACM F | atggcagatctcaattggacaatccccaagacccagaaagc | Constructing plasmid pACM4-YlADH2_02 |
| 20 | YlADH2_02-pACM R | AGCGGTTTCTTTACCAGATTACTTGCTGGTATCGACAAC |  |
| 21 | YlADH2_03-pACM F | atggcagatctcaattggtacaaggacattcccgtcccc | Constructing plasmid pACM4-YlADH2_03 |
| 22 | YlADH2_03-pACM R | AGCGGTTTCTTTACCAGACTACTTGCTGTTATCAACAACG |  |
| 23 | YlADH2_04-pACM F | atggcagatctcaattggtctgctcccgtcatccccaagacc | Constructing plasmid pACM4-YlADH2_04 |
| 24 | YlADH2_04-pACM R | AGCGGTTTCTTTACCAGATTACTTGGAGGTGTCCAGAACG |  |
| 25 | YlADH3-pACM F | atggcagatctcaattggacgcagactcttcccaccacc | Constructing plasmid pACM4-YlADH3 |
| 26 | YlADH3-pACM R | AGCGGTTTCTTTACCAGATCATTTGCTGTTGTCAACAAC |  |
| 27 | YlYPL088W-pACM F | atggcagatctcaattgggtattgatcgaatcagggcatg | Constructing plasmid pACM4-YlYPL088W |
| 28 | YlYPL088W-pACM R | AGCGGTTTCTTTACCAGATTACATCTTGTACTTGACGGGC |  |
| 29 | yqhD-pACM F | atggcagatctcaattggaacaactttaatctgcacaccc | Constructing plasmid pACM4-yqhD |
| 30 | yqhD-pACM R | AGCGGTTTCTTTACCAGATTAGCGGGCGGCTTCGTATATAC |  |
| 31 | adhE-pACM F | atggcagatctcaattgggctgttactaatgtcgctgaac | Constructing plasmid pACM4-adhE |
| 32 | adhE-pACM R | AGCGGTTTCTTTACCAGATTAAGCGGATTTTTTCGCTTTTTTC |  |
| 33 | adhP-pACM F | atggcagatctcaattggaaggctgcagttgttacgaagg | Constructing plasmid pACM4-adhP |
| 34 | adhP-pACM R | AGCGGTTTCTTTACCAGATTAGTGACGGAAATCAATCACC |  |
| 35 | eutG-pACM F | atggcagatctcaattggcaaaatgaattgcagaccgcgc | Constructing plasmid pACM4-eutG |
| 36 | eutG-pACM R | AGCGGTTTCTTTACCAGATTATTGCGCCGCTGCGTACAGG |  |
| 37 | yiaY-pACM F | atggcagatctcaattgggcagcttcaacgttctttattc | Constructing plasmid pACM4-yiaY |
| 38 | yiaY-pACM R | AGCGGTTTCTTTACCAGATTACATCGCTGCGCGATAAATC |  |
| 39 | yahK-pACM F | atggcagatctcaattggaagatcaaagctgttggtgcat | Constructing plasmid pACM4-yahK |
| 40 | yahK-pACM R | AGCGGTTTCTTTACCAGATCAGTCTGTTAGTGTGCGATTATC |  |
| 41 | tyrA(fbr)-pACM4 F | atggcagatctcaattgggttgctgaattgaccgcattacg | Constructing plasmid pACM4-tyrA^fbr^ |
| 42 | tyrA(fbr)-pACM4 R | AGCGGTTTCTTTACCAGATTACTGGCGATTGTCATTCGCC |  |
| 43 | aroG(fbr)-pACM4 F | atggcagatctcaattggaattatcagaacgacgatttacg | Constructing plasmid pACM4-aroG^fbr^ |
| 44 | aroG(fbr)-pACM4 R | AGCGGTTTCTTTACCAGATTACCCGCGACGCGCTTTTACTG |  |
| 45 | gRNA-ΔpheA F | ACGGCGCGAACTGGCCGTCGGTTTTAGAGCTAGAAATAGCAAG | Constructing TargetF-ΔpheA by changing N20 |
| 46 | gRNA-ΔpheA R | CGGCCAGTTCGCGCCGTACTAGCATTATACCTAGGACTG |  |
| 47 | gRNA-ΔtrpE F | ACACAACTGGTGAAAAAGCGGTTTTAGAGCTAGAAATAGCAAG | Constructing TargetF-ΔtrpE by changing N20 |
| 48 | gRNA-ΔtrpE R | TTTTTCACCAGTTGTGTACTAGCATTATACCTAGGACTG |  |
| 49 | gRNA-ΔtyrR F | CTCGATCTACTCGTGCTAAGGTTTTAGAGCTAGAAATAGCAAG | Constructing TargetF-ΔtyrR by changing N20 |
| 50 | gRNA-ΔtyrR R | AGCACGAGTAGATCGAGACTAGCATTATACCTAGGACTG |  |
| 51 | ΔpheA-up F | aatgcatctagatatcctcgagaaacacatctgattaatccac | Constructing *pheA* deletion fragment through over-lapping PCR |
| 52 | ΔpheA-up R | AGTGTTGCCTTTTTGTTATC |  |
| 53 | ΔpheA-down F | aacaaaaaggcaacacttgaaaaggtgccggatgatgtg |  |
| 54 | ΔpheA-down R | TGACCATGATTACGCCGTCGACCATACCAATGGTTTCTGGAGC |  |
| 55 | ΔtrpE-up F | aatgcatctagatatcctcgagaccgtggaaatttccacgccg | Constructing *trpE* deletion fragment through over-lapping PCR |
| 56 | ΔtrpE-up R | TGTTATTCTCTAATTTTGTTC |  |
| 57 | ΔtrpE-down F | aaaattagagaataacatggctgacattctgctgctcg |  |
| 58 | ΔtrpE-down R | TGACCATGATTACGCCGTCGACGAATCCACAAACGCGATCCGC |  |
| 59 | ΔtyrR-up F | aatgcatctagatatcctcgaggtttaattaatcgcatcgccac | Constructing *tyrR* deletion fragment through over-lapping PCR |
| 60 | ΔtyrR-up R | GGGAACCTTCACCTGAAAAAAG |  |
| 61 | ΔtyrR-down F | ttcaggtgaaggttcccgcgcgaatatgcctgatggtgc |  |
| 62 | ΔtyrR-down R | TGACCATGATTACGCCGTCGACCATCCCGCAGGCGGGTAGCAAAG |  |
| 63 | ΔpheA-Cassette F | aaacacatctgattaatccac | Amplifying *pheA* deletion fragment from the vector |
| 64 | ΔpheA-Cassette R | CATACCAATGGTTTCTGGAGC |  |
| 65 | ΔtrpE-Cassette F | accgtggaaatttccacgccg | Amplifying *trpE* deletion fragment from the vector |
| 66 | ΔtrpE-Cassette R | GAATCCACAAACGCGATCCGC |  |
| 67 | ΔtyrR-Cassette F | gtttaattaatcgcatcgccac | Amplifying *tyrR* deletion fragment from the vector |
| 68 | ΔtyrR-Cassette R | CATCCCGCAGGCGGGTAGCAAAG |  |
| 69 | ΔpheA-check F | taaaatttatgacaatgaac | Diagnostic PCR analysis of *pheA* knockout |
| 70 | ΔpheA-check R | ATGATGGTCCGGTGCTGGGG |  |
| 71 | ΔtrpE-check F | tgattgcgccgttctgtctg | Diagnostic PCR analysis of *trpE* knockout |
| 72 | ΔtrpE-check R | AGCGTGTTGGCTGGCTCTAG |  |
| 73 | ΔtyrR-check F | aatcaacgttgatgattgcg | Diagnostic PCR analysis of *tyrR* knockout |
| 74 | ΔtyrR-check R | TAGCATAAACTAGGTGTGACG |  |
| 75 | EcMetK-pCDM4 F | aagaaggagatatacatatggcaaaacacctttttacgtccgag | Constructing plasmid pCDM4-EcMetK |
| 76 | EcMetK-pCDM4 R | AGCGGTTTCTTTACCAGATTACTTCAGACCGGCAGCATCGC |  |
| 77 | BcMetK-pCDM4 F | aagaaggagatatacatatgacaaaaaaacgtcatctgttcac | Constructing plasmid pCDM4-BcMetK |
| 78 | BcMetK-pCDM4 R | AGCGGTTTCTTTACCAGATTATAGACCAGCTTGCTCTTTTAA |  |
| 79 | SaMetK-pCDM4 F | aagaaggagatatacatatgttaaataacaaacgattatttac | Constructing plasmid pCDM4-SaMetK |
| 80 | SaMetK-pCDM4 R | AGCGGTTTCTTTACCAGATTAATATTTTACTGCGTCTTTTAA |  |
| 81 | BsMetK-pCDM4 F | aagaaggagatatacatatgagtaaaaatcgtcgtttatttac | Constructing plasmid pCDM4-BsMetK |
| 82 | BsMetK-pCDM4 R | AGCGGTTTCTTTACCAGATTATTCTCCTAACGCTTCTTTACG |  |
| 83 | ScSAM1-pCDM4 F | aagaaggagatatacatatggccggtacatttttattcacttc | Constructing plasmid pCDM4-ScSAM1 |
| 84 | ScSAM1-pCDM4 R | AGCGGTTTCTTTACCAGATTAGAACTTCAAAGTCTTAGGC |  |
| 85 | ScSAM2-pCDM4 F | aagaaggagatatacatatgtccaagagcaaaactttcttattt | Constructing plasmid pCDM4-ScSAM2 |
| 86 | ScSAM2-pCDM4 R | AGCGGTTTCTTTACCAGATTAAAATTCCAATTTCTTTGGTTT |  |
| 87 | ppc-pCDM4 F | aagaaggagatatacatatgaacgaacaatattccgcattg | Constructing plasmid pCDM4-ppc |
| 88 | ppc-pCDM4 R | AGCGGTTTCTTTACCAGATTAGCCGGTATTACGCATACC |  |
| 89 | metA-pCDM4 F | aagaaggagatatacatatgccgattcgtgtgccggacgagc | Constructing plasmid pCDM4-metA |
| 90 | metA-pCDM4 R | AGCGGTTTCTTTACCAGATTAATCCAGCGTTGGATTCATGTGC |  |
| 91 | metB-pCDM4 F | aagaaggagatatacatatgacgcgtaaacaggccaccatcgcag | Constructing plasmid pCDM4-metB |
| 92 | metB-pCDM4 R | AGCGGTTTCTTTACCAGATTACCCCTTGTTTGCAGCCCGGAAG |  |

Table S3 Plasmids used and developed in this study

| **#** | **Plasmid** | **Characteristics** | **Source or reference** |
| --- | --- | --- | --- |
| 1 | pET28a(PB)N | An ePathBricks vector | Lab stock ^27, 28^ |
| 2 | pACM4 | An ePathBricks vector | Lab stock ^27^ |
| 3 | pCDM4 | An ePathBricks vector | Lab stock ^27^ |
| 4 | pET28a(PB)N-RgTAL | pET28a(PB)N carrying *RgTAL* | This study |
| 5 | pET28a(PB)N-Pc4CL | pET28a(PB)N carrying *Pc4CL* | This study |
| 6 | pET28a(PB)N-LlCCR | pET28a(PB)N carrying *LlCCR* | This study |
| 7 | pET28a(PB)N-ZmCCR | pET28a(PB)N carrying *ZmCCR* | This study |
| 8 | pET28a(PB)N-AtCCR | pET28a(PB)N carrying *AtCCR* | This study |
| 9 | pET28a(PB)N-RgTAL-Pc4CL | pET28a(PB)N carrying *RgTAL* and *Pc4CL* | This study |
| 10 | pET28a(PB)N-RgTAL-Pc4CL-AtCCR | pET28a(PB)N carrying *RgTAL*, *Pc4CL*, and *AtCCR* | This study |
| 11 | pET28a(PB)N-RgTAL-Pc4CL-ZmCCR | pET28a(PB)N carrying *RgTAL*, *Pc4CL*, and *ZmCCR* | This study |
| 12 | pET28a(PB)N-RgTAL-Pc4CL-LlCCR | pET28a(PB)N carrying *RgTAL*, *Pc4CL*, and *LlCCR* | This study |
| 13 | pACM4-ScADH1 | pACM4 carrying *ScADH1* | This study |
| 14 | pACM4-ScADH2 | pACM4 carrying *ScADH2* | This study |
| 15 | pACM4-ScADH3 | pACM4 carrying *ScADH3* | This study |
| 16 | pACM4-ScADH4 | pACM4 carrying *ScADH4* | This study |
| 17 | pACM4-ScADH5 | pACM4 carrying *ScADH5* | This study |
| 18 | pACM4-ScADH6 | pACM4 carrying *ScADH6* | This study |
| 19 | pACM4-ScADH7 | pACM4 carrying *ScADH7* | This study |
| 20 | pACM4-YPL088W | pACM4 carrying *YPL088W* | This study |
| 21 | pACM4-YlADH2_01 | pACM4 carrying *YlADH2_01* | This study |
| 22 | pACM4-YlADH2_02 | pACM4 carrying *YlADH2_02* | This study |
| 23 | pACM4-YlADH2_03 | pACM4 carrying *YlADH2_03* | This study |
| 24 | pACM4-YlADH2_04 | pACM4 carrying *YlADH2_04* | This study |
| 25 | pACM4-YlADH3 | pACM4 carrying *YlADH3* | This study |
| 26 | pACM4-YlYPL088W | pACM4 carrying *YlYPL088W* | This study |
| 27 | pACM4-yqhD | pACM4 carrying *yqhD* | This study |
| 28 | pACM4-adhE | pACM4 carrying *adhE* | This study |
| 29 | pACM4-adhP | pACM4 carrying *adhP* | This study |
| 30 | pACM4-eutG | pACM4 carrying *eutG* | This study |
| 31 | pACM4-yiaY | pACM4 carrying *yiaY* | This study |
| 32 | pACM4-yahK | pACM4 carrying *yahK* | This study |
| 33 | pET28a(PB)N-RgTAL-Pc4CL-AtCCR-ScADH6 | pET28a(PB)N carrying *RgTAL*, *Pc4CL*, *AtCCR*, and *ScADH6* | This study |
| 34 | pACM4-tyrA^fbr^ | pACM4 carrying *tyrA^fbr^* | This study |
| 35 | pACM4-aroG^fbr^ | pACM4 carrying *aroG^fbr^* | This study |
| 36 | pACM4-tyrA^fbr^-aroG^fbr^ | pACM4 carrying *tyrA^fbr^* and *aroG^fbr^* | This study |
| 37 | pCas | CRISPR-Cas9 genome editing tool, expressing Cas9 protein. | From Dr. Yang ^29^ |
| 38 | pTargetF | CRISPR-Cas9 genome editing tool, carrying CRISPR elements. | From Dr. Yang ^29^ |
| 39 | TargetF-ΔtyrR | pTargetF containing N20 specific to *tyrR* | This study |
| 40 | TargetF-ΔpheA | pTargetF containing N20 specific to *pheA* | This study |
| 41 | TargetF-ΔtrpE | pTargetF containing N20 specific to *trpE* | This study |
| 42 | pUC57 | A high copy number cloning vector. | Lab stock |
| 43 | pUC57-ΔtyrR | pUC57 carrying *tyrR* deletion fragment | This study |
| 44 | pUC57-ΔpheA | pUC57 carrying *pheA* deletion fragment | This study |
| 45 | pUC57-ΔtrpE | pUC57 carrying *trpE* deletion fragment | This study |
| 46 | pET28a(PB)N-RgTAL-Pc4CL-AtCCR-PhCFAT | pET28a(PB)N carrying *RgTAL*, *Pc4CL*, *AtCCR*, and *PhCFAT* | This study |
| 47 | pET28a(PB)N-RgTAL-Pc4CL-AtCCR-ScCFAT | pET28a(PB)N carrying *RgTAL*, *Pc4CL*, *AtCCR*, and *ScCFAT* | This study |
| 48 | pET28a(PB)N-RgTAL-Pc4CL-AtCCR-MdAAT1 | pET28a(PB)N carrying *RgTAL*, *Pc4CL*, *AtCCR*, and *MdAAT1* | This study |
| 49 | pET28a(PB)N-RgTAL-Pc4CL-AtCCR-SAAT | pET28a(PB)N carrying *RgTAL*, *Pc4CL*, *AtCCR*, and *SAAT* | This study |
| 50 | pET28a(PB)N-RgTAL-Pc4CL-AtCCR-SlAAT1 | pET28a(PB)N carrying *RgTAL*, *Pc4CL*, *AtCCR*, and *SlAAT1* | This study |
| 51 | pET28a(PB)N-RgTAL-Pc4CL-AtCCR-LtCAAT1 | pET28a(PB)N carrying *RgTAL*, *Pc4CL*, *AtCCR*, and *LtCAAT1* | This study |
| 52 | pET28a(PB)N-RgTAL-Pc4CL-AtCCR-ScCFAT-MdoPhR5 | pET28a(PB)N carrying *RgTAL*, *Pc4CL*, *AtCCR*, *ScCFAT*, and *MdoPhR5* | This study |
| 53 | pET28a(PB)N-RgTAL-Pc4CL-AtCCR-ScCFAT-LtAPS1 | pET28a(PB)N carrying *RgTAL*, *Pc4CL*, *AtCCR*, *ScCFAT*, and *LtAPS1* | This study |
| 54 | pET28a(PB)N-RgTAL-Pc4CL-AtCCR-ScCFAT-OkEGS | pET28a(PB)N carrying *RgTAL*, *Pc4CL*, *AtCCR*, *ScCFAT*, and *OkEGS* | This study |
| 55 | pET28a(PB)N-RgTAL-Pc4CL-AtCCR-PhCFAT-MdoPhR5 | pET28a(PB)N carrying *RgTAL*, *Pc4CL*, *AtCCR*, *PhCFAT*, and *MdoPhR5* | This study |
| 56 | pET28a(PB)N-RgTAL-Pc4CL-AtCCR-MdAAT1-MdoPhR5 | pET28a(PB)N carrying *RgTAL*, *Pc4CL*, *AtCCR*, *MdAAT1*, and *MdoPhR5* | This study |
| 57 | pET28a(PB)N-RgTAL-Pc4CL-AtCCR-SAAT-MdoPhR5 | pET28a(PB)N carrying *RgTAL*, *Pc4CL*, *AtCCR*, *SAAT*, and *MdoPhR5* | This study |
| 58 | pET28a(PB)N-RgTAL-Pc4CL-AtCCR-SlAAT1-MdoPhR5 | pET28a(PB)N carrying *RgTAL*, *Pc4CL*, *AtCCR*, *SlAAT1*, and *MdoPhR5* | This study |
| 59 | pET28a(PB)N-RgTAL-Pc4CL-AtCCR-LtCAAT1-MdoPhR5 | pET28a(PB)N carrying *RgTAL*, *Pc4CL*, *AtCCR*, *LtCAAT1*, and *MdoPhR5* | This study |
| 60 | pET28a(PB)N-RgTAL-Pc4CL-AtCCR-ScCFAT-MdoPhR5-PaAIMT1 | pET28a(PB)N carrying *RgTAL*, *Pc4CL*, *AtCCR*, *ScCFAT*, *MdoPhR5*, and *PaAIMT1* | This study |
| 61 | pET28a(PB)N-RgTAL-Pc4CL-AtCCR-ScCFAT-MdoPhR5-MdoOMT1a | pET28a(PB)N carrying *RgTAL*, *Pc4CL*, *AtCCR*, *ScCFAT*, *MdoPhR5*, and *MdoOMT1a* | This study |
| 62 | pET28a(PB)N-RgTAL-Pc4CL-AtCCR-ScCFAT-MdoPhR5-ObaCVOMT1 | pET28a(PB)N carrying *RgTAL*, *Pc4CL*, *AtCCR*, *ScCFAT*, *MdoPhR5*, and *ObaCVOMT1* | This study |
| 63 | pET28a(PB)N-RgTAL-Pc4CL-AtCCR-ScCFAT-MdoPhR5-ObaEOMT1 | pET28a(PB)N carrying *RgTAL*, *Pc4CL*, *AtCCR*, *ScCFAT*, *MdoPhR5*, and *ObaEOMT1* | This study |
| 64 | pET28a(PB)N-RgTAL-Pc4CL-AtCCR-ScCFAT-MdoPhR5-AtROMT | pET28a(PB)N carrying *RgTAL*, *Pc4CL*, *AtCCR*, *ScCFAT*, *MdoPhR5*, and *AtROMT* | This study |
| 65 | pET28a(PB)N-RgTAL-Pc4CL-AtCCR-ScCFAT-MdoPhR5-OsROMT9 | pET28a(PB)N carrying *RgTAL*, *Pc4CL*, *AtCCR*, *ScCFAT*, *MdoPhR5*, and *OsROMT9* | This study |
| 66 | pET28a(PB)N-RgTAL-Pc4CL-AtCCR-ScCFAT-MdoPhR5-VvROMT | pET28a(PB)N carrying *RgTAL*, *Pc4CL*, *AtCCR*, *ScCFAT*, *MdoPhR5*, and *VvROMT* | This study |
| 67 | pET28a(PB)N-RgTAL-Pc4CL-AtCCR-ScCFAT-MdoPhR5-HlOMT1 | pET28a(PB)N carrying *RgTAL*, *Pc4CL*, *AtCCR*, *ScCFAT*, *MdoPhR5*, and *HlOMT1* | This study |
| 68 | pET28a(PB)N-RgTAL-Pc4CL-AtCCR-ScCFAT-PaAIS1 | pET28a(PB)N carrying *RgTAL*, *Pc4CL*, *AtCCR*, *ScCFAT*, and *PaAIS1* | This study |
| 69 | pET28a(PB)N-RgTAL-Pc4CL-AtCCR-ScCFAT-LtPPS1 | pET28a(PB)N carrying *RgTAL*, *Pc4CL*, *AtCCR*, *ScCFAT*, and *LtPPS1* | This study |
| 70 | pET28a(PB)N-RgTAL-Pc4CL-AtCCR-ScCFAT-PhIGS1 | pET28a(PB)N carrying *RgTAL*, *Pc4CL*, *AtCCR*, *ScCFAT*, and *PhIGS1* | This study |
| 71 | pET28a(PB)N-RgTAL-Pc4CL-AtCCR-ScCFAT-PaAIS1-PaAIMT1 | pET28a(PB)N carrying *RgTAL*, *Pc4CL*, *AtCCR*, *ScCFAT*, *PaAIS1*, and *PaAIMT1* | This study |
| 72 | pET28a(PB)N-RgTAL-Pc4CL-AtCCR-ScCFAT-PaAIS1-MdoOMT1a | pET28a(PB)N carrying *RgTAL*, *Pc4CL*, *AtCCR*, *ScCFAT*, *PaAIS1*, and *MdoOMT1a* | This study |
| 73 | pET28a(PB)N-RgTAL-Pc4CL-AtCCR-ScCFAT-PaAIS1-ObaCVOMT1 | pET28a(PB)N carrying *RgTAL*, *Pc4CL*, *AtCCR*, *ScCFAT*, *PaAIS1*, and *ObaCVOMT1* | This study |
| 74 | pET28a(PB)N-RgTAL-Pc4CL-AtCCR-ScCFAT-PaAIS1-ObaEOMT1 | pET28a(PB)N carrying *RgTAL*, *Pc4CL*, *AtCCR*, *ScCFAT*, *PaAIS1*, and *ObaEOMT1* | This study |
| 75 | pET28a(PB)N-RgTAL-Pc4CL-AtCCR-ScCFAT-PaAIS1-AtROMT | pET28a(PB)N carrying *RgTAL*, *Pc4CL*, *AtCCR*, *ScCFAT*, *PaAIS1*, and *AtROMT* | This study |
| 76 | pET28a(PB)N-RgTAL-Pc4CL-AtCCR-ScCFAT-PaAIS1-OsROMT9 | pET28a(PB)N carrying *RgTAL*, *Pc4CL*, *AtCCR*, *ScCFAT*, *PaAIS1*, and *OsROMT9* | This study |
| 77 | pET28a(PB)N-RgTAL-Pc4CL-AtCCR-ScCFAT-PaAIS1-VvROMT | pET28a(PB)N carrying *RgTAL*, *Pc4CL*, *AtCCR*, *ScCFAT*, *PaAIS1*, and *VvROMT* | This study |
| 78 | pET28a(PB)N-RgTAL-Pc4CL-AtCCR-ScCFAT-PaAIS1-HlOMT1 | pET28a(PB)N carrying *RgTAL*, *Pc4CL*, *AtCCR*, *ScCFAT*, *PaAIS1*, and *HlOMT1* | This study |
| 79 | pET28a(PB)N-RgTAL-Pc4CL-AtCCR-ScCFAT-LtPPS1-PaAIMT1 | pET28a(PB)N carrying *RgTAL*, *Pc4CL*, *AtCCR*, *ScCFAT*, *LtPPS1*, and *PaAIMT1* | This study |
| 80 | pET28a(PB)N-RgTAL-Pc4CL-AtCCR-ScCFAT-LtPPS1-MdoOMT1a | pET28a(PB)N carrying *RgTAL*, *Pc4CL*, *AtCCR*, *ScCFAT*, *LtPPS1*, and *MdoOMT1a* | This study |
| 81 | pET28a(PB)N-RgTAL-Pc4CL-AtCCR-ScCFAT-LtPPS1-ObaCVOMT1 | pET28a(PB)N carrying *RgTAL*, *Pc4CL*, *AtCCR*, *ScCFAT*, *LtPPS1*, and *ObaCVOMT1* | This study |
| 82 | pET28a(PB)N-RgTAL-Pc4CL-AtCCR-ScCFAT-LtPPS1-ObaEOMT1 | pET28a(PB)N carrying *RgTAL*, *Pc4CL*, *AtCCR*, *ScCFAT*, *LtPPS1*, and *ObaEOMT1* | This study |
| 83 | pET28a(PB)N-RgTAL-Pc4CL-AtCCR-ScCFAT-LtPPS1-AtROMT | pET28a(PB)N carrying *RgTAL*, *Pc4CL*, *AtCCR*, *ScCFAT*, *LtPPS1*, and *AtROMT* | This study |
| 84 | pET28a(PB)N-RgTAL-Pc4CL-AtCCR-ScCFAT-LtPPS1-OsROMT9 | pET28a(PB)N carrying *RgTAL*, *Pc4CL*, *AtCCR*, *ScCFAT*, *LtPPS1*, and *OsROMT9* | This study |
| 85 | pET28a(PB)N-RgTAL-Pc4CL-AtCCR-ScCFAT-LtPPS1-VvROMT | pET28a(PB)N carrying *RgTAL*, *Pc4CL*, *AtCCR*, *ScCFAT*, *LtPPS1*, and *VvROMT* | This study |
| 86 | pET28a(PB)N-RgTAL-Pc4CL-AtCCR-ScCFAT-LtPPS1-HlOMT1 | pET28a(PB)N carrying *RgTAL*, *Pc4CL*, *AtCCR*, *ScCFAT*, *LtPPS1*, and *HlOMT1* | This study |
| 87 | pET28a(PB)N-RgTAL-Pc4CL-AtCCR-ScCFAT-PhIGS1-PaAIMT1 | pET28a(PB)N carrying *RgTAL*, *Pc4CL*, *AtCCR*, *ScCFAT*, *PhIGS1*, and *PaAIMT1* | This study |
| 88 | pET28a(PB)N-RgTAL-Pc4CL-AtCCR-ScCFAT-PhIGS1-MdoOMT1a | pET28a(PB)N carrying *RgTAL*, *Pc4CL*, *AtCCR*, *ScCFAT*, *PhIGS1*, and *MdoOMT1a* | This study |
| 89 | pET28a(PB)N-RgTAL-Pc4CL-AtCCR-ScCFAT-PhIGS1-ObaCVOMT1 | pET28a(PB)N carrying *RgTAL*, *Pc4CL*, *AtCCR*, *ScCFAT*, *PhIGS1*, and *ObaCVOMT1* | This study |
| 90 | pET28a(PB)N-RgTAL-Pc4CL-AtCCR-ScCFAT-PhIGS1-ObaEOMT1 | pET28a(PB)N carrying *RgTAL*, *Pc4CL*, *AtCCR*, *ScCFAT*, *PhIGS1*, and *ObaEOMT1* | This study |
| 91 | pET28a(PB)N-RgTAL-Pc4CL-AtCCR-ScCFAT-PhIGS1-AtROMT | pET28a(PB)N carrying *RgTAL*, *Pc4CL*, *AtCCR*, *ScCFAT*, *PhIGS1*, and *AtROMT* | This study |
| 92 | pET28a(PB)N-RgTAL-Pc4CL-AtCCR-ScCFAT-PhIGS1-OsROMT9 | pET28a(PB)N carrying *RgTAL*, *Pc4CL*, *AtCCR*, *ScCFAT*, *PhIGS1*, and *OsROMT9* | This study |
| 93 | pET28a(PB)N-RgTAL-Pc4CL-AtCCR-ScCFAT-PhIGS1-VvROMT | pET28a(PB)N carrying *RgTAL*, *Pc4CL*, *AtCCR*, *ScCFAT*, *PhIGS1*, and *VvROMT* | This study |
| 94 | pET28a(PB)N-RgTAL-Pc4CL-AtCCR-ScCFAT-PhIGS1-HlOMT1 | pET28a(PB)N carrying *RgTAL*, *Pc4CL*, *AtCCR*, *ScCFAT*, *PhIGS1*, and *HlOMT1* | This study |
| 95 | pACM4-tyrA^fbr^-aroG^fbr^-RgTAL | pACM4 carrying *tyrA^fbr^*, *aroG^fbr^*, and *RgTAL* | This study |
| 96 | pACM4-tyrA^fbr^-aroG^fbr^-4CL | pACM4 carrying *tyrA^fbr^*, *aroG^fbr^*, and *4CL* | This study |
| 97 | pACM4-tyrA^fbr^-aroG^fbr^-AtCCR | pACM4 carrying *tyrA^fbr^*, *aroG^fbr^*, and *AtCCR* | This study |
| 98 | pACM4-tyrA^fbr^-aroG^fbr^-ScADH6 | pACM4 carrying *tyrA^fbr^*, *aroG^fbr^*, and *ScADH6* | This study |
| 99 | pACM4-tyrA^fbr^-aroG^fbr^-ScCFAT | pACM4 carrying *tyrA^fbr^*, *aroG^fbr^*, and *ScCFAT* | This study |
| 100 | pACM4-tyrA^fbr^-aroG^fbr^-PaAIS1 | pACM4 carrying *tyrA^fbr^*, *aroG^fbr^*, and *PaAIS1* | This study |
| 101 | pACM4-tyrA^fbr^-aroG^fbr^-ObaCVOMT1 | pACM4 carrying *tyrA^fbr^*, *aroG^fbr^*, and *ObaCVOMT1* | This study |
| 102 | pACM4-tyrA^fbr^-aroG^fbr^-ObaCVOMT1-ObaCVOMT1 | pACM4 carrying *tyrA^fbr^*, *aroG^fbr^*, and 2 copies *ObaCVOMT1* | This study |
| 103 | pACM4-tyrA^fbr^-aroG^fbr^-RgTAL | pACM4 carrying *tyrA^fbr^*, *aroG^fbr^*, and *RgTAL* | This study |
| 104 | pACM4-tyrA^fbr^-aroG^fbr^-4CL | pACM4 carrying *tyrA^fbr^*, *aroG^fbr^*, and *4CL* | This study |
| 105 | pACM4-tyrA^fbr^-aroG^fbr^-AtCCR1 | pACM4 carrying *tyrA^fbr^*, *aroG^fbr^*, and *AtCCR1* | This study |
| 106 | pACM4-tyrA^fbr^-aroG^fbr^-ScADH6 | pACM4 carrying *tyrA^fbr^*, *aroG^fbr^*, and *ScADH6* | This study |
| 107 | pACM4-tyrA^fbr^-aroG^fbr^-ScCFAT | pACM4 carrying *tyrA^fbr^*, *aroG^fbr^*, and *ScCFAT* | This study |
| 108 | pACM4-tyrA^fbr^-aroG^fbr^-PaAIS1 | pACM4 carrying *tyrA^fbr^*, *aroG^fbr^*, and *PaAIS1* | This study |
| 109 | pACM4-tyrA^fbr^-aroG^fbr^-ObaEOMT1 | pACM4 carrying *tyrA^fbr^*, *aroG^fbr^*, and *ObaEOMT1* | This study |
| 110 | pACM4-tyrA^fbr^-aroG^fbr^-ObaEOMT1-ObaEOMT1 | pACM4 carrying *tyrA^fbr^*, *aroG^fbr^*, and 2 copies *ObaEOMT1* | This study |
| 111 | pCDM4-EcMetK | pCDM4 carrying *EcMetK* | This study |
| 112 | pCDM4-BcMetK | pCDM4 carrying *BcMetK* | This study |
| 113 | pCDM4-SaMetK | pCDM4 carrying *SaMetK* | This study |
| 114 | pCDM4-BsMetK | pCDM4 carrying *BsMetK* | This study |
| 115 | pCDM4-ScSAM1 | pCDM4 carrying *ScSAM1* | This study |
| 116 | pCDM4-ScSAM2 | pCDM4 carrying *ScSAM2* | This study |
| 117 | pCDM4-ppc-BcMetK | pCDM4 carrying *ppc*, and *BcMetK* | This study |
| 118 | pCDM4-metA-BcMetK | pCDM4 carrying *metA*, and *BcMetK* | This study |
| 119 | pCDM4-metB-BcMetK | pCDM4 carrying *metB*, and *BcMetK* | This study |

Table S4 Strains used and developed in this study

| **#** | **Strain** | **Description** | **Source or reference** |
| --- | --- | --- | --- |
| 1 | *E. coli* JM109 | Molecular biology, plasmid construction and propagation. | Lab stock |
| 2 | *E. coli* BL21(DE3) | Gene expression, pathway analysis, and production of estragole, anethole, and their derivatives. | Lab stock |
| 3 | BL21/pET | *E. coli* BL21(DE3) containing pET28a(PB)N | This study |
| 4 | BL21/pACM | *E. coli* BL21(DE3) containing pACM4 | This study |
| 5 | BL21/pET-pACM | *E. coli* BL21(DE3) containing pET28a(PB)N and pACM4 | This study |
| 6 | BL21/T4 | *E. coli* BL21(DE3) containing pET28a(PB)N-RgTAL-Pc4CL | This study |
| 7 | BL21/AtCCR | *E. coli* BL21(DE3) containing pET28a(PB)N-AtCCR | This study |
| 8 | BL21/ZmCCR | *E. coli* BL21(DE3) containing pET28a(PB)N-ZmCCR | This study |
| 9 | BL21/LlCCR | *E. coli* BL21(DE3) containing pET28a(PB)N-LlCCR | This study |
| 10 | CAD-01 | *E. coli* BL21(DE3) containing pET28a(PB)N-RgTAL-Pc4CL-AtCCR | This study |
| 11 | CAD-02 | *E. coli* BL21(DE3) containing pET28a(PB)N-RgTAL-Pc4CL-ZmCCR | This study |
| 12 | CAD-03 | *E. coli* BL21(DE3) containing pET28a(PB)N-RgTAL-Pc4CL-LlCCR | This study |
| 13 | BL21/ScADH1 | *E. coli* BL21(DE3) containing pACM4-ScADH1 | This study |
| 14 | BL21/ScADH2 | *E. coli* BL21(DE3) containing pACM4-ScADH2 | This study |
| 15 | BL21/ScADH3 | *E. coli* BL21(DE3) containing pACM4-ScADH3 | This study |
| 16 | BL21/ScADH4 | *E. coli* BL21(DE3) containing pACM4-ScADH4 | This study |
| 17 | BL21/ScADH5 | *E. coli* BL21(DE3) containing pACM4-ScADH5 | This study |
| 18 | BL21/ScADH6 | *E. coli* BL21(DE3) containing pACM4-ScADH6 | This study |
| 19 | BL21/ScADH7 | *E. coli* BL21(DE3) containing pACM4-ScADH7 | This study |
| 20 | BL21/YPL088W | *E. coli* BL21(DE3) containing pACM4-YPL088W | This study |
| 21 | BL21/YlADH2_01 | *E. coli* BL21(DE3) containing pACM4-YlADH2_01 | This study |
| 22 | BL21/YlADH2_02 | *E. coli* BL21(DE3) containing pACM4-YlADH2_02 | This study |
| 23 | BL21/YlADH2_03 | *E. coli* BL21(DE3) containing pACM4-YlADH2_03 | This study |
| 24 | BL21/YlADH2_04 | *E. coli* BL21(DE3) containing pACM4-YlADH2_04 | This study |
| 25 | BL21/YlADH3 | *E. coli* BL21(DE3) containing pACM4-YlADH3 | This study |
| 26 | BL21/YlYPL088W | *E. coli* BL21(DE3) containing pACM4-YlYPL088W | This study |
| 27 | BL21/yqhD | *E. coli* BL21(DE3) containing pACM4-yqhD | This study |
| 28 | BL21/adhE | *E. coli* BL21(DE3) containing pACM4-adhE | This study |
| 29 | BL21/adhP | *E. coli* BL21(DE3) containing pACM4-adhP | This study |
| 30 | BL21/eutG | *E. coli* BL21(DE3) containing pACM4-eutG | This study |
| 31 | BL21/yiaY | *E. coli* BL21(DE3) containing pACM4-yiaY | This study |
| 32 | BL21/yahK | *E. coli* BL21(DE3) containing pACM4-yahK | This study |
| 33 | BL21/aroG^fbr^ | *E. coli* BL21(DE3) containing pACM4-aroG^frb^ | This study |
| 34 | BL21/tyrA^fbr^ | *E. coli* BL21(DE3) containing pACM4-tyrA^fbr^ | This study |
| 35 | CAL-01 | *E. coli* BL21(DE3) containing pET28a(PB)N-RgTAL-Pc4CL-AtCCR and pACM4-ScADH1 | This study |
| 36 | CAL-02 | *E. coli* BL21(DE3) containing pET28a(PB)N-RgTAL-Pc4CL-AtCCR and pACM4-ScADH2 | This study |
| 37 | CAL-03 | *E. coli* BL21(DE3) containing pET28a(PB)N-RgTAL-Pc4CL-AtCCR and pACM4-ScADH3 | This study |
| 38 | CAL-04 | *E. coli* BL21(DE3) containing pET28a(PB)N-RgTAL-Pc4CL-AtCCR and pACM4-ScADH4 | This study |
| 39 | CAL-05 | *E. coli* BL21(DE3) containing pET28a(PB)N-RgTAL-Pc4CL-AtCCR and pACM4-ScADH5 | This study |
| 40 | CAL-06 | *E. coli* BL21(DE3) containing pET28a(PB)N-RgTAL-Pc4CL-AtCCR and pACM4-ScADH6 | This study |
| 41 | CAL-07 | *E. coli* BL21(DE3) containing pET28a(PB)N-RgTAL-Pc4CL-AtCCR and pACM4-ScADH7 | This study |
| 42 | CAL-08 | *E. coli* BL21(DE3) containing pET28a(PB)N-RgTAL-Pc4CL-AtCCR and pACM4-YPL088W | This study |
| 43 | CAL-09 | *E. coli* BL21(DE3) containing pET28a(PB)N-RgTAL-Pc4CL-AtCCR and pACM4-YlADH2_01 | This study |
| 44 | CAL-10 | *E. coli* BL21(DE3) containing pET28a(PB)N-RgTAL-Pc4CL-AtCCR and pACM4-YlADH2_02 | This study |
| 45 | CAL-11 | *E. coli* BL21(DE3) containing pET28a(PB)N-RgTAL-Pc4CL-AtCCR and pACM4-YlADH2_03 | This study |
| 46 | CAL-12 | *E. coli* BL21(DE3) containing pET28a(PB)N-RgTAL-Pc4CL-AtCCR and pACM4-YlADH2_04 | This study |
| 47 | CAL-13 | *E. coli* BL21(DE3) containing pET28a(PB)N-RgTAL-Pc4CL-AtCCR and pACM4-YlADH3 | This study |
| 48 | CAL-14 | *E. coli* BL21(DE3) containing pET28a(PB)N-RgTAL-Pc4CL-AtCCR and pACM4-YlYPL088W | This study |
| 49 | CAL-15 | *E. coli* BL21(DE3) containing pET28a(PB)N-RgTAL-Pc4CL-AtCCR and pACM4-yqhD | This study |
| 50 | CAL-16 | *E. coli* BL21(DE3) containing pET28a(PB)N-RgTAL-Pc4CL-AtCCR and pACM4-adhE | This study |
| 51 | CAL-17 | *E. coli* BL21(DE3) containing pET28a(PB)N-RgTAL-Pc4CL-AtCCR and pACM4-adhP | This study |
| 52 | CAL-18 | *E. coli* BL21(DE3) containing pET28a(PB)N-RgTAL-Pc4CL-AtCCR and pACM4-eutG | This study |
| 53 | CAL-19 | *E. coli* BL21(DE3) containing pET28a(PB)N-RgTAL-Pc4CL-AtCCR and pACM4-yiaY | This study |
| 54 | CAL-20 | *E. coli* BL21(DE3) containing pET28a(PB)N-RgTAL-Pc4CL-AtCCR and pACM4-yahK | This study |
| 55 | CAD-01/pACM | *E. coli* BL21(DE3) containing pET28a(PB)N-RgTAL-Pc4CL-AtCCR and empty pACM4 | This study |
| 56 | CAL-21 | *E. coli* BL21(DE3) containing pET28a(PB)N-RgTAL-Pc4CL-AtCCR-ScADH6 | This study |
| 57 | CAL-22 | *E. coli* BL21(DE3) containing pET28a(PB)N-RgTAL-Pc4CL-AtCCR and pACM4-aroG^fbr^ | This study |
| 58 | CAL-23 | *E. coli* BL21(DE3) containing pET28a(PB)N-RgTAL-Pc4CL-AtCCR and pACM4-tyrA^fbr^ | This study |
| 59 | CAL-24 | *E. coli* BL21(DE3) containing pET28a(PB)N-RgTAL-Pc4CL-AtCCR and pACM4-tyrA^fbr^-aroG^fbr^ | This study |
| 60 | ΔA | *E. coli* BL21(DE3) removing *pheA* gene | This study |
| 61 | ΔE | *E. coli* BL21(DE3) removing *trpE* gene | This study |
| 62 | ΔR | *E. coli* BL21(DE3) removing *tyrR* gene | This study |
| 63 | ΔAE | *E. coli* BL21(DE3) removing *pheA* and *trpE* genes | This study |
| 64 | ΔRA | *E. coli* BL21(DE3) removing *tyrR* and *pheA* genes | This study |
| 65 | ΔRE | *E. coli* BL21(DE3) removing *tyrR* and *trpE* genes | This study |
| 66 | ΔRAE | *E. coli* BL21(DE3) removing *tyrR*, *pheA*, and *trpE* genes | This study |
| 67 | CAL-25 | ΔR containing pET28a(PB)N-RgTAL-Pc4CL-AtCCR and pACM4-tyrA^fbr^-aroG^fbr^ | This study |
| 68 | CAL-26 | ΔRA containing pET28a(PB)N-RgTAL-Pc4CL-AtCCR and pACM4-tyrA^fbr^-aroG^fbr^ | This study |
| 69 | CAL-27 | ΔRE containing pET28a(PB)N-RgTAL-Pc4CL-AtCCR and pACM4-tyrA^fbr^-aroG^fbr^ | This study |
| 70 | CAL-28 | ΔRAE containing pET28a(PB)N-RgTAL-Pc4CL-AtCCR and pACM4-tyrA^fbr^-aroG^fbr^ | This study |
| 71 | CAL-29 | ΔA containing pET28a(PB)N-RgTAL-Pc4CL-AtCCR and pACM4-tyrA^fbr^-aroG^fbr^ | This study |
| 72 | CAL-30 | ΔE containing pET28a(PB)N-RgTAL-Pc4CL-AtCCR and pACM4-tyrA^fbr^-aroG^fbr^ | This study |
| 73 | CAL-31 | ΔAE containing pET28a(PB)N-RgTAL-Pc4CL-AtCCR and pACM4-tyrA^fbr^-aroG^fbr^ | This study |
| 74 | CAT-01 | ΔRAE containing pET28a(PB)N-RgTAL-Pc4CL-AtCCR-PhCFAT and pACM4-tyrA^fbr^-aroG^fbr^ | This study |
| 75 | CAT-02 | ΔRAE containing pET28a(PB)N-RgTAL-Pc4CL-AtCCR-ScCFAT and pACM4-tyrA^fbr^-aroG^fbr^ | This study |
| 76 | CAT-03 | ΔRAE containing pET28a(PB)N-RgTAL-Pc4CL-AtCCR-MdAAT1 and pACM4-tyrA^fbr^-aroG^fbr^ | This study |
| 77 | CAT-04 | ΔRAE containing pET28a(PB)N-RgTAL-Pc4CL-AtCCR-SAAT and pACM4-tyrA^fbr^-aroG^fbr^ | This study |
| 78 | CAT-05 | ΔRAE containing pET28a(PB)N-RgTAL-Pc4CL-AtCCR-SlAAT1 and pACM4-tyrA^fbr^-aroG^fbr^ | This study |
| 79 | CAT-06 | ΔRAE containing pET28a(PB)N-RgTAL-Pc4CL-AtCCR-LtCAAT1 and pACM4-tyrA^fbr^-aroG^fbr^ | This study |
| 80 | CHA-01 | ΔRAE containing pET28a(PB)N-RgTAL-Pc4CL-AtCCR-ScCFAT-MdoPhR5 and pACM4-tyrA^fbr^-aroG^fbr^ | This study |
| 81 | CHA-02 | ΔRAE containing pET28a(PB)N-RgTAL-Pc4CL-AtCCR-ScCFAT-LtAPS1 and pACM4-tyrA^fbr^-aroG^fbr^ | This study |
| 82 | CHA-03 | ΔRAE containing pET28a(PB)N-RgTAL-Pc4CL-AtCCR-ScCFAT-OkEGS and pACM4-tyrA^fbr^-aroG^fbr^ | This study |
| 83 | CHA-04 | ΔRAE containing pET28a(PB)N-RgTAL-Pc4CL-AtCCR-PhCFAT-MdoPhR5 and pACM4-tyrA^fbr^-aroG^fbr^ | This study |
| 84 | CHA-05 | ΔRAE containing pET28a(PB)N-RgTAL-Pc4CL-AtCCR-MdAAT1-MdoPhR5 and pACM4-tyrA^fbr^-aroG^fbr^ | This study |
| 85 | CHA-06 | ΔRAE containing pET28a(PB)N-RgTAL-Pc4CL-AtCCR-SAAT-MdoPhR5 and pACM4-tyrA^fbr^-aroG^fbr^ | This study |
| 86 | CHA-07 | ΔRAE containing pET28a(PB)N-RgTAL-Pc4CL-AtCCR-SlAAT1-MdoPhR5 and pACM4-tyrA^fbr^-aroG^fbr^ | This study |
| 87 | CHA-08 | ΔRAE containing pET28a(PB)N-RgTAL-Pc4CL-AtCCR-LtCAAT1-MdoPhR5 and pACM4-tyrA^fbr^-aroG^fbr^ | This study |
| 88 | EST-01 | ΔRAE containing pET28a(PB)N-RgTAL-Pc4CL-AtCCR-ScCFAT-MdoPhR5-PaAIMT1 and pACM4-tyrA^fbr^-aroG^fbr^ | This study |
| 89 | EST-02 | ΔRAE containing pET28a(PB)N-RgTAL-Pc4CL-AtCCR-ScCFAT-MdoPhR5-MdoOMT1a and pACM4-tyrA^fbr^-aroG^fbr^ | This study |
| 90 | EST-03 | ΔRAE containing pET28a(PB)N-RgTAL-Pc4CL-AtCCR-ScCFAT-MdoPhR5-ObaCVOMT1 and pACM4-tyrA^fbr^-aroG^fbr^ | This study |
| 91 | EST-04 | ΔRAE containing pET28a(PB)N-RgTAL-Pc4CL-AtCCR-ScCFAT-MdoPhR5-ObaEOMT1 and pACM4-tyrA^fbr^-aroG^fbr^ | This study |
| 92 | EST-05 | ΔRAE containing pET28a(PB)N-RgTAL-Pc4CL-AtCCR-ScCFAT-MdoPhR5-AtROMT and pACM4-tyrA^fbr^-aroG^fbr^ | This study |
| 93 | EST-06 | ΔRAE containing pET28a(PB)N-RgTAL-Pc4CL-AtCCR-ScCFAT-MdoPhR5-OsROMT9 and pACM4-tyrA^fbr^-aroG^fbr^ | This study |
| 94 | EST-07 | ΔRAE containing pET28a(PB)N-RgTAL-Pc4CL-AtCCR-ScCFAT-MdoPhR5-VvROMT and pACM4-tyrA^fbr^-aroG^fbr^ | This study |
| 95 | EST-08 | ΔRAE containing pET28a(PB)N-RgTAL-Pc4CL-AtCCR-ScCFAT-MdoPhR5-HlOMT1 and pACM4-tyrA^fbr^-aroG^fbr^ | This study |
| 96 | EST-09 | ΔRAE containing pET28a(PB)N-RgTAL-Pc4CL-AtCCR-ScCFAT-MdoPhR5-ObaCVOMT1 and pACM4-tyrA^fbr^-aroG^fbr^-RgTAL | This study |
| 97 | EST-10 | ΔRAE containing pET28a(PB)N-RgTAL-Pc4CL-AtCCR-ScCFAT-MdoPhR5-ObaCVOMT1 and pACM4-tyrA^fbr^-aroG^fbr^-4CL | This study |
| 98 | EST-11 | ΔRAE containing pET28a(PB)N-RgTAL-Pc4CL-AtCCR-ScCFAT-MdoPhR5-ObaCVOMT1 and pACM4-tyrA^fbr^-aroG^fbr^-AtCCR | This study |
| 99 | EST-12 | ΔRAE containing pET28a(PB)N-RgTAL-Pc4CL-AtCCR-ScCFAT-MdoPhR5-ObaCVOMT1 and pACM4-tyrA^fbr^-aroG^fbr^-ScADH6 | This study |
| 100 | EST-13 | ΔRAE containing pET28a(PB)N-RgTAL-Pc4CL-AtCCR-ScCFAT-MdoPhR5-ObaCVOMT1 and pACM4-tyrA^fbr^-aroG^fbr^-ScCFAT | This study |
| 101 | EST-14 | ΔRAE containing pET28a(PB)N-RgTAL-Pc4CL-AtCCR-ScCFAT-MdoPhR5-ObaCVOMT1 and pACM4-tyrA^fbr^-aroG^fbr^-PaAIS1 | This study |
| 102 | EST-15 | ΔRAE containing pET28a(PB)N-RgTAL-Pc4CL-AtCCR-ScCFAT-MdoPhR5-ObaCVOMT1 and pACM4-tyrA^fbr^-aroG^fbr^-ObaCVOMT1 | This study |
| 103 | EST-16 | ΔRAE containing pET28a(PB)N-RgTAL-Pc4CL-AtCCR-ScCFAT-MdoPhR5-ObaCVOMT1 and pACM4-tyrA^fbr^-aroG^fbr^-ObaCVOMT1-ObaCVOMT1 | This study |
| 104 | EST-17 | ΔRAE containing pET28a(PB)N-RgTAL-Pc4CL-AtCCR-ScCFAT-MdoPhR5-ObaCVOMT1, pACM4-tyrA^fbr^-aroG^fbr^-ObaCVOMT1-ObaCVOMT1, and pCDM4-EcMetK |  |
| 105 | EST-18 | ΔRAE containing pET28a(PB)N-RgTAL-Pc4CL-AtCCR-ScCFAT-MdoPhR5-ObaCVOMT1, pACM4-tyrA^fbr^-aroG^fbr^-ObaCVOMT1-ObaCVOMT1, and pCDM4-BcMetK |  |
| 106 | EST-19 | ΔRAE containing pET28a(PB)N-RgTAL-Pc4CL-AtCCR-ScCFAT-MdoPhR5-ObaCVOMT1, pACM4-tyrA^fbr^-aroG^fbr^-ObaCVOMT1-ObaCVOMT1, and pCDM4-SaMetK |  |
| 107 | EST-20 | ΔRAE containing pET28a(PB)N-RgTAL-Pc4CL-AtCCR-ScCFAT-MdoPhR5-ObaCVOMT1, pACM4-tyrA^fbr^-aroG^fbr^-ObaCVOMT1-ObaCVOMT1, and pCDM4-BsMetK |  |
| 108 | EST-21 | ΔRAE containing pET28a(PB)N-RgTAL-Pc4CL-AtCCR-ScCFAT-MdoPhR5-ObaCVOMT1, pACM4-tyrA^fbr^-aroG^fbr^-ObaCVOMT1-ObaCVOMT1, and pCDM4-ScSAM1 |  |
| 109 | EST-22 | ΔRAE containing pET28a(PB)N-RgTAL-Pc4CL-AtCCR-ScCFAT-MdoPhR5-ObaCVOMT1, pACM4-tyrA^fbr^-aroG^fbr^-ObaCVOMT1-ObaCVOMT1, and pCDM4-ScSAM2 |  |
| 110 | EST-23 | ΔRAE containing pET28a(PB)N-RgTAL-Pc4CL-AtCCR-ScCFAT-MdoPhR5-ObaCVOMT1, pACM4-tyrA^fbr^-aroG^fbr^-ObaCVOMT1-ObaCVOMT1, and pCDM4-ppc-BcMetK |  |
| 111 | EST-24 | ΔRAE containing pET28a(PB)N-RgTAL-Pc4CL-AtCCR-ScCFAT-MdoPhR5-ObaCVOMT1, pACM4-tyrA^fbr^-aroG^fbr^-ObaCVOMT1-ObaCVOMT1, and pCDM4-metA-BcMetK |  |
| 112 | EST-25 | ΔRAE containing pET28a(PB)N-RgTAL-Pc4CL-AtCCR-ScCFAT-MdoPhR5-ObaCVOMT1, pACM4-tyrA^fbr^-aroG^fbr^-ObaCVOMT1-ObaCVOMT1, and pCDM4-metB-BcMetK |  |
| 113 | ISC-01 | ΔRAE containing pET28a(PB)N-RgTAL-Pc4CL-AtCCR-ScCFAT-PaAIS1 and pACM4-tyrA^fbr^-aroG^fbr^ | This study |
| 114 | ISC-02 | ΔRAE containing pET28a(PB)N-RgTAL-Pc4CL-AtCCR-ScCFAT-LtPPS1 and pACM4-tyrA^fbr^-aroG^fbr^ | This study |
| 115 | ISC-03 | ΔRAE containing pET28a(PB)N-RgTAL-Pc4CL-AtCCR-ScCFAT-PhIGS1 and pACM4-tyrA^fbr^-aroG^fbr^ | This study |
| 116 | ANT-01 | ΔRAE containing pET28a(PB)N-RgTAL-Pc4CL-AtCCR-ScCFAT-PaAIS1-PaAIMT1 and pACM4-tyrA^fbr^-aroG^fbr^ | This study |
| 117 | ANT-02 | ΔRAE containing pET28a(PB)N-RgTAL-Pc4CL-AtCCR-ScCFAT-PaAIS1-MdoOMT1a and pACM4-tyrA^fbr^-aroG^fbr^ | This study |
| 118 | ANT-03 | ΔRAE containing pET28a(PB)N-RgTAL-Pc4CL-AtCCR-ScCFAT-PaAIS1-ObaCVOMT1 and pACM4-tyrA^fbr^-aroG^fbr^ | This study |
| 119 | ANT-04 | ΔRAE containing pET28a(PB)N-RgTAL-Pc4CL-AtCCR-ScCFAT-PaAIS1-ObaEOMT1 and pACM4-tyrA^fbr^-aroG^fbr^ | This study |
| 120 | ANT-05 | ΔRAE containing pET28a(PB)N-RgTAL-Pc4CL-AtCCR-ScCFAT-PaAIS1-AtROMT and pACM4-tyrA^fbr^-aroG^fbr^ | This study |
| 121 | ANT-06 | ΔRAE containing pET28a(PB)N-RgTAL-Pc4CL-AtCCR-ScCFAT-PaAIS1-OsROMT9 and pACM4-tyrA^fbr^-aroG^fbr^ | This study |
| 122 | ANT-07 | ΔRAE containing pET28a(PB)N-RgTAL-Pc4CL-AtCCR-ScCFAT-PaAIS1-VvROMT and pACM4-tyrA^fbr^-aroG^fbr^ | This study |
| 123 | ANT-08 | ΔRAE containing pET28a(PB)N-RgTAL-Pc4CL-AtCCR-ScCFAT-PaAIS1-HlOMT1 and pACM4-tyrA^fbr^-aroG^fbr^ | This study |
| 124 | ANT-09 | ΔRAE containing pET28a(PB)N-RgTAL-Pc4CL-AtCCR-ScCFAT-LtPPS1-PaAIMT1 and pACM4-tyrA^fbr^-aroG^fbr^ | This study |
| 125 | ANT-10 | ΔRAE containing pET28a(PB)N-RgTAL-Pc4CL-AtCCR-ScCFAT-LtPPS1-MdoOMT1a and pACM4-tyrA^fbr^-aroG^fbr^ | This study |
| 126 | ANT-11 | ΔRAE containing pET28a(PB)N-RgTAL-Pc4CL-AtCCR-ScCFAT-LtPPS1-ObaCVOMT1 and pACM4-tyrA^fbr^-aroG^fbr^ | This study |
| 127 | ANT-12 | ΔRAE containing pET28a(PB)N-RgTAL-Pc4CL-AtCCR-ScCFAT-LtPPS1-ObaEOMT1 and pACM4-tyrA^fbr^-aroG^fbr^ | This study |
| 128 | ANT-13 | ΔRAE containing pET28a(PB)N-RgTAL-Pc4CL-AtCCR-ScCFAT-LtPPS1-AtROMT and pACM4-tyrA^fbr^-aroG^fbr^ | This study |
| 129 | ANT-14 | ΔRAE containing pET28a(PB)N-RgTAL-Pc4CL-AtCCR-ScCFAT-LtPPS1-OsROMT9 and pACM4-tyrA^fbr^-aroG^fbr^ | This study |
| 130 | ANT-15 | ΔRAE containing pET28a(PB)N-RgTAL-Pc4CL-AtCCR-ScCFAT-LtPPS1-VvROMT and pACM4-tyrA^fbr^-aroG^fbr^ | This study |
| 131 | ANT-16 | ΔRAE containing pET28a(PB)N-RgTAL-Pc4CL-AtCCR-ScCFAT-LtPPS1-HlOMT1 and pACM4-tyrA^fbr^-aroG^fbr^ | This study |
| 132 | ANT-17 | ΔRAE containing pET28a(PB)N-RgTAL-Pc4CL-AtCCR-ScCFAT-PhIGS1-PaAIMT1 and pACM4-tyrA^fbr^-aroG^fbr^ | This study |
| 133 | ANT-18 | ΔRAE containing pET28a(PB)N-RgTAL-Pc4CL-AtCCR-ScCFAT-PhIGS1-MdoOMT1a and pACM4-tyrA^fbr^-aroG^fbr^ | This study |
| 134 | ANT-19 | ΔRAE containing pET28a(PB)N-RgTAL-Pc4CL-AtCCR-ScCFAT-PhIGS1-ObaCVOMT1 and pACM4-tyrA^fbr^-aroG^fbr^ | This study |
| 135 | ANT-20 | ΔRAE containing pET28a(PB)N-RgTAL-Pc4CL-AtCCR-ScCFAT-PhIGS1-ObaEOMT1 and pACM4-tyrA^fbr^-aroG^fbr^ | This study |
| 136 | ANT-21 | ΔRAE containing pET28a(PB)N-RgTAL-Pc4CL-AtCCR-ScCFAT-PhIGS1-AtROMT and pACM4-tyrA^fbr^-aroG^fbr^ | This study |
| 137 | ANT-22 | ΔRAE containing pET28a(PB)N-RgTAL-Pc4CL-AtCCR-ScCFAT-PhIGS1-OsROMT9 and pACM4-tyrA^fbr^-aroG^fbr^ | This study |
| 138 | ANT-23 | ΔRAE containing pET28a(PB)N-RgTAL-Pc4CL-AtCCR-ScCFAT-PhIGS1-VvROMT and pACM4-tyrA^fbr^-aroG^fbr^ | This study |
| 139 | ANT-24 | ΔRAE containing pET28a(PB)N-RgTAL-Pc4CL-AtCCR-ScCFAT-PhIGS1-HlOMT1 and pACM4-tyrA^fbr^-aroG^fbr^ | This study |
| 140 | ANT-25 | ΔRAE containing pET28a(PB)N-RgTAL-Pc4CL-AtCCR-ScCFAT-PaAIS1-ObaEOMT1 and pACM4-tyrA^fbr^-aroG^fbr^-RgTAL | This study |
| 141 | ANT-26 | ΔRAE containing pET28a(PB)N-RgTAL-Pc4CL-AtCCR-ScCFAT-PaAIS1-ObaEOMT1 and pACM4-tyrA^fbr^-aroG^fbr^-4CL | This study |
| 142 | ANT-27 | ΔRAE containing pET28a(PB)N-RgTAL-Pc4CL-AtCCR-ScCFAT-PaAIS1-ObaEOMT1 and pACM4-tyrA^fbr^-aroG^fbr^-AtCCR1 | This study |
| 143 | ANT-28 | ΔRAE containing pET28a(PB)N-RgTAL-Pc4CL-AtCCR-ScCFAT-PaAIS1-ObaEOMT1 and pACM4-tyrA^fbr^-aroG^fbr^-ScADH6 | This study |
| 144 | ANT-29 | ΔRAE containing pET28a(PB)N-RgTAL-Pc4CL-AtCCR-ScCFAT-PaAIS1-ObaEOMT1 and pACM4-tyrA^fbr^-aroG^fbr^-ScCFAT | This study |
| 145 | ANT-30 | ΔRAE containing pET28a(PB)N-RgTAL-Pc4CL-AtCCR-ScCFAT-PaAIS1-ObaEOMT1 and pACM4-tyrA^fbr^-aroG^fbr^-PaAIS1 | This study |
| 146 | ANT-31 | ΔRAE containing pET28a(PB)N-RgTAL-Pc4CL-AtCCR-ScCFAT-PaAIS1-ObaEOMT1 and pACM4-tyrA^fbr^-aroG^fbr^-ObaEOMT1 | This study |
| 147 | ANT-32 | ΔRAE containing pET28a(PB)N-RgTAL-Pc4CL-AtCCR-ScCFAT-PaAIS1-ObaEOMT1 and pACM4-tyrA^fbr^-aroG^fbr^-ObaEOMT1-ObaEOMT1 | This study |
| 148 | ANT-33 | ΔRAE containing pET28a(PB)N-RgTAL-Pc4CL-AtCCR-ScCFAT-PaAIS1-ObaEOMT1, pACM4-tyrA^fbr^-aroG^fbr^-ObaEOMT1-ObaEOMT1, and pCDM4-EcMetK | This study |
| 149 | ANT-34 | ΔRAE containing pET28a(PB)N-RgTAL-Pc4CL-AtCCR-ScCFAT-PaAIS1-ObaEOMT1, pACM4-tyrA^fbr^-aroG^fbr^-ObaEOMT1-ObaEOMT1, and pCDM4-BcMetK | This study |
| 150 | ANT-35 | ΔRAE containing pET28a(PB)N-RgTAL-Pc4CL-AtCCR-ScCFAT-PaAIS1-ObaEOMT1, pACM4-tyrA^fbr^-aroG^fbr^-ObaEOMT1-ObaEOMT1, and pCDM4-SaMetK | This study |
| 151 | ANT-36 | ΔRAE containing pET28a(PB)N-RgTAL-Pc4CL-AtCCR-ScCFAT-PaAIS1-ObaEOMT1, pACM4-tyrA^fbr^-aroG^fbr^-ObaEOMT1-ObaEOMT1, and pCDM4-BsMetK | This study |
| 152 | ANT-37 | ΔRAE containing pET28a(PB)N-RgTAL-Pc4CL-AtCCR-ScCFAT-PaAIS1-ObaEOMT1, pACM4-tyrA^fbr^-aroG^fbr^-ObaEOMT1-ObaEOMT1, and pCDM4-ScSAM1 | This study |
| 153 | ANT-38 | ΔRAE containing pET28a(PB)N-RgTAL-Pc4CL-AtCCR-ScCFAT-PaAIS1-ObaEOMT1, pACM4-tyrA^fbr^-aroG^fbr^-ObaEOMT1-ObaEOMT1, and pCDM4-ScSAM2 | This study |
| 154 | ANT-39 | ΔRAE containing pET28a(PB)N-RgTAL-Pc4CL-AtCCR-ScCFAT-PaAIS1-ObaEOMT1, pACM4-tyrA^fbr^-aroG^fbr^-ObaEOMT1-ObaEOMT1, and pCDM4-ppc-BcMetK | This study |
| 155 | ANT-40 | ΔRAE containing pET28a(PB)N-RgTAL-Pc4CL-AtCCR-ScCFAT-PaAIS1-ObaEOMT1, pACM4-tyrA^fbr^-aroG^fbr^-ObaEOMT1-ObaEOMT1, and pCDM4-metA-BcMetK | This study |
| 156 | ANT-41 | ΔRAE containing pET28a(PB)N-RgTAL-Pc4CL-AtCCR-ScCFAT-PaAIS1-ObaEOMT1, pACM4-tyrA^fbr^-aroG^fbr^-ObaEOMT1-ObaEOMT1, and pCDM4-metB-BcMetK | This study |
